# Epigenetic Rescue of Disuse Muscle Atrophy by Histone Deacetylase Inhibitor Trichostatin A

**DOI:** 10.64898/2026.10.30.763257

**Authors:** Kanita Hrustanovic, Scott Burlingham, Katie Sikes, Ryan Foslien, Kenna Thomas, Lauren Monroe, Benjamin C Gadmoski, Soham Ghosh

## Abstract

Mechanical unloading rapidly induces skeletal muscle atrophy and functional deterioration, yet how the loss of mechanical input influences nuclear architecture, chromatin organization, and associated transcriptional programs remains poorly understood. Here, we investigated whether pharmacological inhibition of histone deacetylases (HDACs) using Trichostatin A (TSA) could attenuate the consequences of continued mechanical unloading in a murine hindlimb suspension model. Following 14 days of unloading, mice exhibited impaired locomotor activity, altered muscle architecture, and widespread transcriptional reprogramming characterized by increased expression of atrophy associated genes, inflammatory mediators, and extracellular matrix remodeling pathways, alongside suppression of mitochondrial metabolic programs. Super resolution imaging further revealed irregular Lamin A/C organization and disruption of discrete H3K9me3 associated heterochromatin foci. Systemic TSA treatment significantly attenuated the decline in locomotor activity, accompanied by trends toward reduced body weight loss, and preservation of muscle morphology. Transcriptomic analysis demonstrated a substantial shift toward the baseline expression profile, with high-dose TSA attenuating FOXO associated catabolic programs, inflammatory, and extracellular matrix-related signatures, while enhancing mitochondrial oxidative phosphorylation pathways. Quantitative gene level and pathway analyses further revealed dose dependent and selective reversal of unloading associated transcriptional changes. Notably, TSA treatment also partially preserved nuclear lamina organization and the focal distribution of H3K9me3 associated heterochromatin. Despite these muscle specific benefits, TSA did not prevent trabecular bone loss over the 14-day unloading period, although several muscle-expressed genes potentially involved in muscle to bone communication were altered by treatment. Together, these findings demonstrate that HDAC inhibition counteracts multiple functional and molecular consequences of skeletal muscle unloading and is associated with partial preservation of nuclear and heterochromatin organization. This study supports a link between mechanical unloading, nuclear architectural remodeling, and HDAC sensitive transcriptional regulation, providing a foundation for epigenetic strategies to counter disuse induced muscle deterioration.

## INTRODUCTION

Skeletal muscle disuse atrophy, characterized by the progressive loss of muscle mass and function following reduced mechanical loading, is a significant clinical concern affecting individuals across different ages and health conditions^1–3^. It commonly develops during prolonged bed rest, limb immobilization following injury or surgery, and neurological conditions that restrict normal muscle activity. Although disuse atrophy is mechanistically distinct from age-associated sarcopenia and cancer cachexia, these conditions can coexist and further accelerate muscle deterioration^4,5^. Reduced mechanical loading is also a major contributor to skeletal muscle loss during spaceflight, highlighting the fundamental importance of mechanical stimuli in maintaining muscle homeostasis muscle^6,7^. The consequences of disuse develop rapidly, particularly in older adults, with even short periods of bed rest producing substantial reductions in muscle mass, strength, and functional capacity^8^. These changes can delay rehabilitation, increase the risk of falls and mobility impairment, and contribute to prolonged dependence on clinical care. Current management relies primarily on rehabilitation and exercise-based interventions, often supported by nutritional strategies^9^. However, these approaches may be difficult to implement in patients with severe mobility restrictions or prolonged immobilization, and no established pharmacological treatment is specifically approved to prevent disuse-induced muscle atrophy. Identifying molecular targets that can preserve muscle function during periods of reduced mechanical loading therefore remains an important therapeutic goal.

Maintenance of skeletal muscle mass depends on the coordinated regulation of protein synthesis and degradation, mitochondrial metabolism, and tissue remodeling, all of which respond to mechanical activity^10–12^. Under physiological loading conditions, anabolic signaling through pathways such as IGF-1/PI3K/Akt/mTORC1 contributes to protein synthesis and muscle growth. Mechanical unloading disrupts this balance by reducing anabolic activity and activating catabolic programs, including FOXO-associated transcriptional regulation of the ubiquitin-proteasome and autophagy-lysosome systems^13,14^. Increased expression of muscle-specific ubiquitin ligases, particularly MuRF1 (*Trim63*) and Atrogin-1 (*Fbxo32*), promotes the degradation of muscle proteins and contributes to progressive muscle wasting. Beyond protein turnover, unloading induces changes in mitochondrial metabolism, inflammatory signaling, extracellular matrix organization, and muscle regeneration, collectively altering the structural and functional state of skeletal muscle^1–3^. Although these pathways have been extensively investigated, how the loss of mechanical input coordinates such widespread transcriptional responses remains incompletely understood.

Mechanical forces regulate cellular behavior not only through conventional cytoplasmic signaling pathways but also through their transmission to the cell nucleus^15–19^. Mechanical signals originating at the extracellular matrix and sarcolemma can be transmitted through the cytoskeleton and the linker of nucleoskeleton and cytoskeleton (LINC) complex to the nuclear lamina. Nuclear envelope proteins, including Lamin A/C, emerin, and associated structural components, contribute to the mechanical stability of the nucleus and influence its interactions with chromatin. Changes in nuclear deformation and mechanical coupling can consequently alter chromatin organization and gene regulation, providing a potential connection between the mechanical environment and cellular transcriptional responses. Emerging evidence indicates that mechanical forces influence the spatial organization of heterochromatin and the regulation of histone modifications. In our previous work, we demonstrated that nuclear deformation contributes to the organization of H3K9me3-associated heterochromatin during cardiac development and that disruption of mechanical coupling alters this organization and associated gene expression^20^. Epigenetic regulation has also been implicated in skeletal muscle adaptation, including the persistence of transcriptional responses following previous mechanical loading, a phenomenon often described as muscle epigenetic memory^21,22^. However, the relationship between reduced mechanical loading, nuclear architecture, and epigenetic regulation during skeletal muscle disuse remains poorly defined.

Histone deacetylases (HDACs) have emerged as important regulators of skeletal muscle homeostasis and atrophy. These enzymes remove acetyl groups from histones and several non-histone proteins, influencing chromatin-associated transcriptional regulation as well as the activity of signaling proteins. HDAC4 has been implicated in the induction of atrophy-associated genes following denervation, including *Gadd45a* and myogenic regulatory factors programs^23^, while class I HDAC activity has been shown to promote FOXO-dependent catabolic signaling during muscle disuse^24^. Pharmacological inhibition of HDACs has consequently attracted interest as a potential strategy to limit muscle wasting. Notably, previous studies demonstrated that Trichostatin A (TSA), a broad-spectrum HDAC inhibitor, partially attenuates muscle fiber atrophy and suppresses unloading-associated MuRF1 expression in the rat soleus muscle following hindlimb suspension^25^. These findings establish that HDAC inhibition can influence the muscle atrophy response. However, it remains unclear whether HDAC inhibition broadly reverses the transcriptional consequences of mechanical unloading, which biological programs are preferentially responsive to treatment, and whether such responses are accompanied by preservation of nuclear architecture and heterochromatin organization. In particular, the potential connection between HDAC-sensitive transcriptional regulation and mechanically induced changes in nuclear structure has received comparatively limited attention in the context of skeletal muscle disuse.

In the present study, we investigated the effects of TSA on skeletal muscle function, transcriptional regulation, and nuclear organization using a murine hindlimb suspension (HLS) model over 14 days. We hypothesized that mechanical unloading alters nuclear lamina and heterochromatin organization alongside widespread transcriptional changes and that HDAC inhibition can attenuate these responses. We evaluated the effects of two TSA doses on unloading-associated functional decline and muscle morphology using locomotor assessment, ultrasound imaging, and histological analysis. Bulk RNA sequencing was performed to characterize the global transcriptional consequences of unloading, identify HDAC-sensitive biological pathways, and quantify the extent and dose dependence of transcriptional rescue. Super-resolution immunofluorescence imaging was used to examine changes in Lamin A/C organization and the distribution of H3K9me3-associated heterochromatin. Given the functional coupling between skeletal muscle and bone, we also assessed femoral architecture and mechanical properties and examined muscle-derived genes implicated in muscle–bone crosstalk. By integrating functional, transcriptomic, and nuclear structural analyses, this study seeks to establish a broader understanding of the HDAC-sensitive response to mechanical unloading and evaluate the potential of epigenetic intervention to mitigate disuse-associated skeletal muscle deterioration.

## MATERIALS AND METHODS

### Animal care and study approval

All animal studies were conducted upon approval of the IACUC at Colorado State University. 16-week-old female C57BL/6J mice were purchased from the Jackson Laboratory. The animals were housed in a pathogen-free environment care facility at Colorado State University, with a 12-hour dark/light cycle and unrestricted access to food and water. Food and enrichment were placed on the bed of rodent cages for ease of access during unloading. Two mice were housed per cage for all experiments. Mice were observed by trained veterinary specialists for any signs of distress and for immediate corrective action if the harness was loose.

### Animal models and treatment groups

An adapted Morey-Holton Hindlimb Suspension (HLS) system was used to simulate disuse induced muscle atrophy in mice, which was originally designed for rats (Fig. 1A). Mice were suspended for 14 days before euthanasia for downstream functional assays, tissue harvesting followed by all assays. For this study, two concentrations of drug were used. Low dose TSA: 1.5 mg/Kg of mouse body weight (Trichostatin A, Ready Made Solution, T1952, Millipore Sigma), and high dose: 8 mg/Kg of mouse body weight in a 200µl injection volume (Trichostatin A, T8552, Millipore Sigma). Harnesses for hindlimb were made of 10-gauge aluminum wire and 1/8” ID x 3/16” OD medical-grade silicone tubing, wrapped in fleece fabric for added cushion. A small carabiner hook was used to attach the harness to suspension rod. Rodent’s front limbs were fully weight bearing while hindlimbs were fully suspended. Mice were checked daily for harness fitment and monitoring. The following five experimental groups were used: NS (No Suspension, mice freely move inside the cage), control (HLS), DMSO (vehicle for drug + HLS), HLS + low TSA, and HLS + high TSA. TSA and DMSO groups received weekly subcutaneous injections at the back of the neck at day 0 (right before suspension) and day 7. For toxicity studies, a separate cohort of normally loaded (non-suspended) mice received subcutaneous injections of low and high-dose TSA in the same dosing manner. Mice were under anesthesia (isofluorane) during the harness installation and drug administration. Mice were euthanized using carbon di-oxide overdose or cardiac poke followed by cervical dislocation.

**Figure 1.**
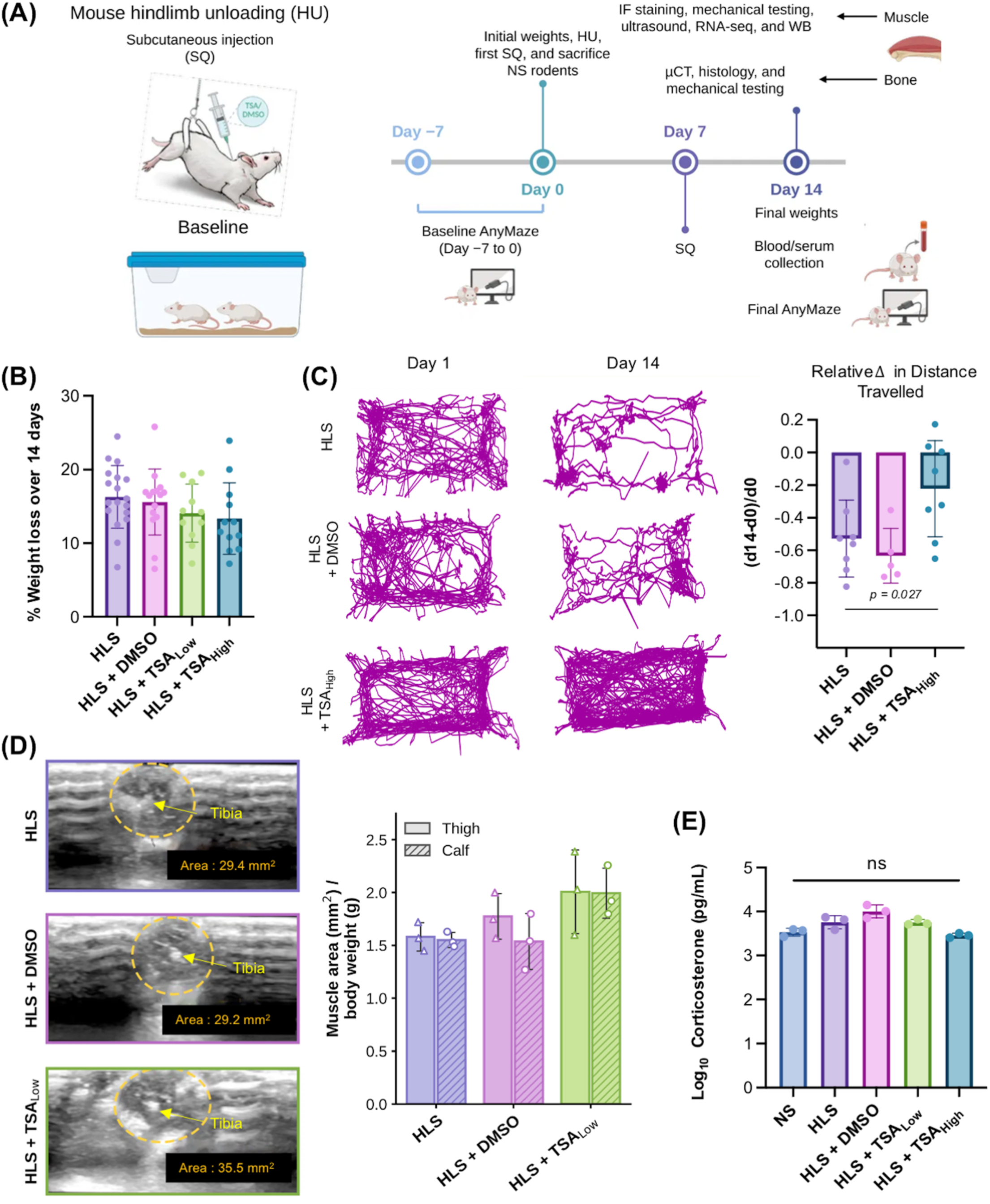
Trichostatin A (TSA) treatment attenuates locomotor decline and is associated with reduced body-weight loss following 14 days of hindlimb suspension. **(A)** Schematic of experimental design. Mice underwent 14 days of harness-based hindlimb suspension (HLS) to induce mechanical unloading. Five experimental groups were included: no suspension (NS), HLS, HLS + DMSO (vehicle control), HLS + low TSA (1.5 mg/kg body weight), and HLS + high TSA (8 mg/kg body weight). Mice receiving DMSO or TSA were administered by two subcutaneous injectionson days 0 and 7. Body weights and locomotor activity were assessed before and after suspension, followed by blood and tissue collection for downstream analyses. **(B)** Percentage of body-weight loss over the 14-day experimental period in HLS, HLS + DMSO, HLS + low TSA, and HLS + high TSA groups. TSA-treated mice exhibited a trend toward reduced body-weight loss, with the lowest average loss observed in the HLS + high TSA group. Error bars represent mean ± SD. **(C)** Representative ANY-maze movement trajectories before and after hindlimb suspension for HLS, HLS + DMSO, and HLS + high TSA mice. Quantification of the relative change in distance traveled, calculated as (day 14 − day 0)/day 0, demonstrated a significantly smaller decline in locomotor activity following high-dose TSA treatment compared with HLS (*p* = 0.027) and HLS + DMSO.. Error bars represent mean ± SD, groups were compared by Kruskal–Wallis test followed by Dunn’s multiple comparisons test. **(D)** Representative transverse ultrasound images of the calf muscles after 14 days of hindlimb suspension, with outlined regions indicating the measured muscle areas. Quantification of thigh and calf muscle cross-sectional area, normalized to terminal body weight, showed a trend toward larger muscle areas in HLS + low TSA mice compared with HLS and HLS + DMSO. **(E)** Serum corticosterone concentrations measured by ELISA at the 14-day endpoint. No statistically significant differences were detected among NS, HLS, HLS + DMSO, HLS + low TSA, and HLS + high TSA groups.

### Assessment of body weight and functionality

Mice body weight was measured on days 0 and 14 and percent change in weight loss was calculated (day 14-day 0/day 0 × 100). Mice mobility was assessed before and after hindlimb suspension using the ANY-maze video tracking system to quantify their mobility related functional decline, including changes in distance traveled and freezing episodes along with other parameters. All parameters were calculated and normalized to baseline: % Change = (Post-HLS value − mean pre-HLS value)/(mean pre-HLS value) × 100. All mice underwent two acclimation and two baseline sessions (averaged results used), with 10-minute tracking recordings per session. A final post-HLS assessment was conducted 14 days after suspension, preceded by a 1-hour re-acclimation period without harnesses to minimize stress-induced behavioral artifacts. Each rodent was alone in their cage with no hut present during this process. Functional assessments were conducted at the same time of day to minimize variability in the results.

### Corticosterone ELISA assay and serum panel for toxicity

Serum was isolated from whole blood collected via cardiac puncture immediately after mice were euthanized and stored at –80°C until further analysis. Corticosterone levels were quantified using Enzo ELISA kit (ADI-901-097) according to the manufacturer’s instructions. A microplate reader was used to measure absorbance at 450 nm, and corticosterone concentrations were calculated based on a standard curve.

### Ultrasound imaging

Transverse ultrasound (US) scans of the cadaveric hindlimbs (calf and quadriceps muscles) were acquired using a high-frequency linear probe (GE Healthcare, Logic E R8. Probe: GE L8-18i-RS Ultrasound Transducer). Cross-sectional area (CSA) measurements based on the US images were performed by an unbiased Veterinary Technician using Horos (v3.6.6, Horos Project), using the femur, tibia, and fibula as consistent anatomical landmarks to ensure reproducibility. To account for individual variations in atrophy severity, CSA values were normalized to each animal’s final weight after HLS.

### RNA sequencing

Total RNA was extracted from flash-frozen gastrocnemius muscle using TRIzol™ reagent with bead homogenization, followed by purification on RNeasy columns (Qiagen). RNA integrity was assessed using an Agilent Bioanalyzer, and samples meeting quality criteria (RIN ≥ [7]) were shipped on dry ice to Novogene (Sacramento, CA) for sequencing. Novogene performed additional quality control, including quantification (NanoDrop and Agilent), purity assessment (NanoDrop and agarose gel electrophoresis), and integrity confirmation (Agilent Bioanalyzer). Poly(A)-enriched libraries were prepared and sequenced on an Illumina [NovaSeq X Plus] platform to generate 150 bp paired-end reads at a target depth of 40 million reads per sample.

### RNA sequencing analysis

Raw reads were quality-filtered with [fastp], aligned to the mouse reference genome ([GRCm39/mm39]) using [HISAT2], and gene-level counts were generated with [featureCounts]. Differential expression analysis was performed in R (version 4.4.3) using DESeq2. Each HLS group (HLS, HLS + DMSO, HLS + TSA_Low, HLS + TSA_High) was compared with non-suspended (NS) controls, and each treated HLS group was compared with untreated HLS, using the Wald test. P-values were adjusted by the Benjamini–Hochberg method, and genes with padj < 0.05 and |log2 fold change| ≥ 1 were considered differentially expressed. Gene symbols were assigned from Ensembl IDs using org.Mm.eg.db. For heatmaps and clustering, counts were transformed with DESeq2’s variance-stabilizing transformation (VST) and z-scored per gene. For individual gene plots, normalized counts were expressed relative to the NS group mean and shown as mean ± SEM, with significance determined by DESeq2 adjusted p-values.

Over-representation analysis of up- and downregulated genes was performed with clusterProfiler for GO (Biological Process, Cellular Component, Molecular Function) and KEGG pathways, using all tested genes as the background (BH-adjusted p < 0.05, q < 0.20); redundant GO terms were collapsed using a semantic similarity cutoff of 0.7. Gene set enrichment analysis (GSEA) was performed on genes ranked by the DESeq2 Wald statistic using GO Biological Process terms and MSigDB Hallmark gene sets (msigdbr, mouse), with BH-adjusted p < 0.05 considered significant.

To assess transcriptional rescue by TSA, analyses were restricted to HLS-altered genes (HLS vs NS, padj < 0.05, |log2FC| ≥ 1). The fraction of the HLS effect reversed by treatment was calculated as 1 − |log2FC(HLS vs NS) + log2FC(TSA + HLS vs HLS)| / |log2FC(HLS vs NS)|, and genes were considered significantly rescued when the treatment effect opposed the HLS effect and was significant relative to HLS (padj < 0.05). Global reversal was estimated by regressing each treatment’s log2FC versus NS on the HLS-vs-NS log2FC through the origin, with percent reversal defined as 100 × (1 − slope). Co-regulated gene modules were identified by k-means clustering (k = 6, 50 random starts) of z-scored group-mean VST values for HLS- and TSA-responsive genes, followed by GO enrichment of each module. Finally, a curated panel of muscle-expressed genes involved in muscle-bone communication (myokines, bone-signal receptors, Wnt/β-catenin, TGF-β/BMP, RANKL/OPG, and osteogenic genes) was examined across comparisons, and bone-related GO pathways were tested by GSEA.

### Mechanical characterization of bone

Fourteen days after unloading, murine femurs were harvested, flash-frozen in liquid nitrogen, and stored at -80°C until further use. Femurs designated for µCT analysis were thawed and immediately fixed in 10% neutral buffered formalin (for 48 hours) before scanning at 10 µm isotropic resolution using a Scanco Medical µCT 80 system. Standard quantification of bone mineral density (BMD) and bone volume fraction (BV/TV) was performed for both the distal trabecular and mid-diaphyseal cortical regions using the manufacturer’s software (Scanco Medical Evaluation software). Concurrently, femurs selected for mechanical testing were maintained in phosphate-buffered saline (PBS) until further testing. Uniaxial 3-point bending tests were performed on an MTS 858 MiniBionix II mechanical testing system equipped with a 100 N load cell, with femurs positioned on supports and a load applied at the mid-diaphysis at a constant displacement rate of 0.01mm/s oscillating between 0.05-5 N for 5 preconditioning cycles, followed by a single ramp to failure at the same displacement rate. Subsequently, bone stiffness was calculated from the linear elastic region of the load-displacement curves from the ramp to failure portion of the test.

### Histology of bone and muscle

Bones with adjacent quadriceps muscle were fixed in 10% neutral-buffered formalin. Following fixation, samples were decalcified in 10% EDTA for ∼14 days on a rocker. Tissues were then embedded in paraffin blocks in both the sagittal and transverse orientations and sectioned at a thickness of 5 µm using a rotary microtome. Sections were transferred to microscope slides and prepared for histological staining by deparaffinization and rehydration through a series of graded alcohols and distilled water. Sections were stained with hematoxylin and eosin (H&E) and Masson’s trichrome following an established staining protocol. Images of histological sections were captured using a Zeiss 980 microscope.

### Immunofluorescence (IF) staining and imaging

Murine calf muscles were harvested, fixed in 4% paraformaldehyde (PFA) for 24 hours at 4°C, then thoroughly washed in PBS and embedded in low-melting point agarose (4%). Tissues were sliced into 150 µm thick sections using a Vibratome followed by IF staining. After antigen retrieval, tissues were permeabilized, blocked for non-specific binding sites. Sections were incubated for two nights at 4°C with primary antibodies targeting H3K9me3 (Invitrogen, 1:500) and Lamin A/C (Invitrogen, 1:500), followed by washes and 1-hour incubation with fluorophore-conjugated secondary antibodies (Alexa Fluor 488 or Alexa Fluor 594, Invitrogen). DAPI (Invitrogen, catalog #: D21490) was included for nuclear counterstaining. Images were acquired on a Zeiss confocal laser scanning microscope with AiryScan 2 using consistent settings using the super-resolution mode.

## RESULTS

### Trichostatin A treatment attenuates functional decline following hindlimb suspension

To determine whether pharmacological inhibition of histone deacetylases (HDACs) could mitigate the effects of mechanical unloading on skeletal muscle, we used a murine hindlimb suspension (HLS) model and administered the pan-HDAC inhibitor Trichostatin A (TSA) over a 14-day period (Fig. 1A). Mice were assigned to five experimental groups: no suspension (NS), HLS, HLS + DMSO (vehicle), HLS + low TSA (1.5 mg/kg), and HLS + high TSA (8 mg/kg). TSA or DMSO was administered through weekly subcutaneous injections, and the effects of treatment were evaluated through changes in body weight, locomotor activity, and gross muscle morphology, as measured by ultrasound.

Following 14 days of unloading, mice in the HLS and HLS + DMSO groups exhibited substantial body-weight loss (Fig. 1B). Both TSA-treated groups showed a lower average percentage of weight loss, with the greatest reduction observed in the HLS + high TSA group. Although these differences did not establish a statistically significant treatment effect, the trend suggested that TSA treatment may mitigate the loss of body mass associated with prolonged mechanical unloading.

We next assessed whether TSA treatment could preserve locomotor function following hindlimb unloading. Mice underwent movement tracking using the ANY-maze system before and after the 14-day suspension period. Representative movement trajectories revealed a marked reduction in voluntary locomotor activity in HLS and HLS + DMSO mice after unloading, whereas mice receiving HLS + high TSA retained more extensive movement patterns (Fig. 1C). Quantitative analysis confirmed that the relative decline in distance traveled was smaller in the HLS + high TSA group compared with both HLS and HLS + DMSO (*p* = 0.027). Additional locomotor parameters supported this functional improvement. The increase in freezing episodes was attenuated in HLS + high TSA mice compared with HLS mice (*p* = 0.05), while changes in mean speed and the number of body rotations showed similar trends toward improved locomotor activity (Fig. S1A). Additional measurements from mice receiving low-dose TSA are presented in Fig. S1B. Together, these findings indicate that high-dose TSA treatment partially preserves locomotor activity despite prolonged mechanical unloading.

To assess whether these functional improvements were accompanied by changes in muscle morphology, we performed ultrasound imaging of the hindlimb muscles after 14 days of unloading. Representative transverse ultrasound images revealed differences in muscle dimensions among treatment groups (Fig. 1D). Quantification of muscle cross-sectional area, normalized to terminal body weight, showed a trend toward larger quadricep and gastrocnemius muscle areas in the HLS + low TSA group compared with HLS and HLS + DMSO. Additional measurements of muscle area from longitudinal and transverse ultrasound images are presented in Fig. S1D. Although these measurements suggest a possible preservation of muscle dimensions following TSA treatment, the observed differences were variable and did not establish a significant treatment effect.

Finally, we measured circulating corticosterone, a physiological marker of stress, to determine whether the experimental procedures or drug treatment were associated with altered stress hormone levels. Serum corticosterone concentrations measured at the 14-day endpoint were comparable across NS, HLS, HLS + DMSO, HLS + low TSA, and HLS + high TSA groups, with no statistically significant differences detected (Fig. 1E). These findings suggest that neither hindlimb suspension nor TSA treatment produced a detectable difference in circulating corticosterone at the time of tissue collection.

Collectively, these results demonstrate that systemic HDAC inhibition with TSA attenuates the decline in locomotor activity following 14 days of hindlimb suspension. The accompanying trends in body-weight loss and muscle dimensions further suggest a potential protective effect against unloading-induced deterioration, motivating a more detailed investigation of the tissue-level and molecular mechanisms underlying TSA treatment.

### Assessment of systemic effects and potential toxicity following TSA treatment

To determine whether TSA administration was associated with systemic toxicity, we evaluated changes in body weight, relative organ weights, and serum biochemical markers following low-dose (1.5 mg/kg) and high-dose (8 mg/kg) TSA treatment. These assessments were performed in non-suspended mice receiving TSA, while serum biochemical measurements were additionally performed in HLS + low TSA and HLS + high TSA mice (Fig. 2).

**Figure 2.**
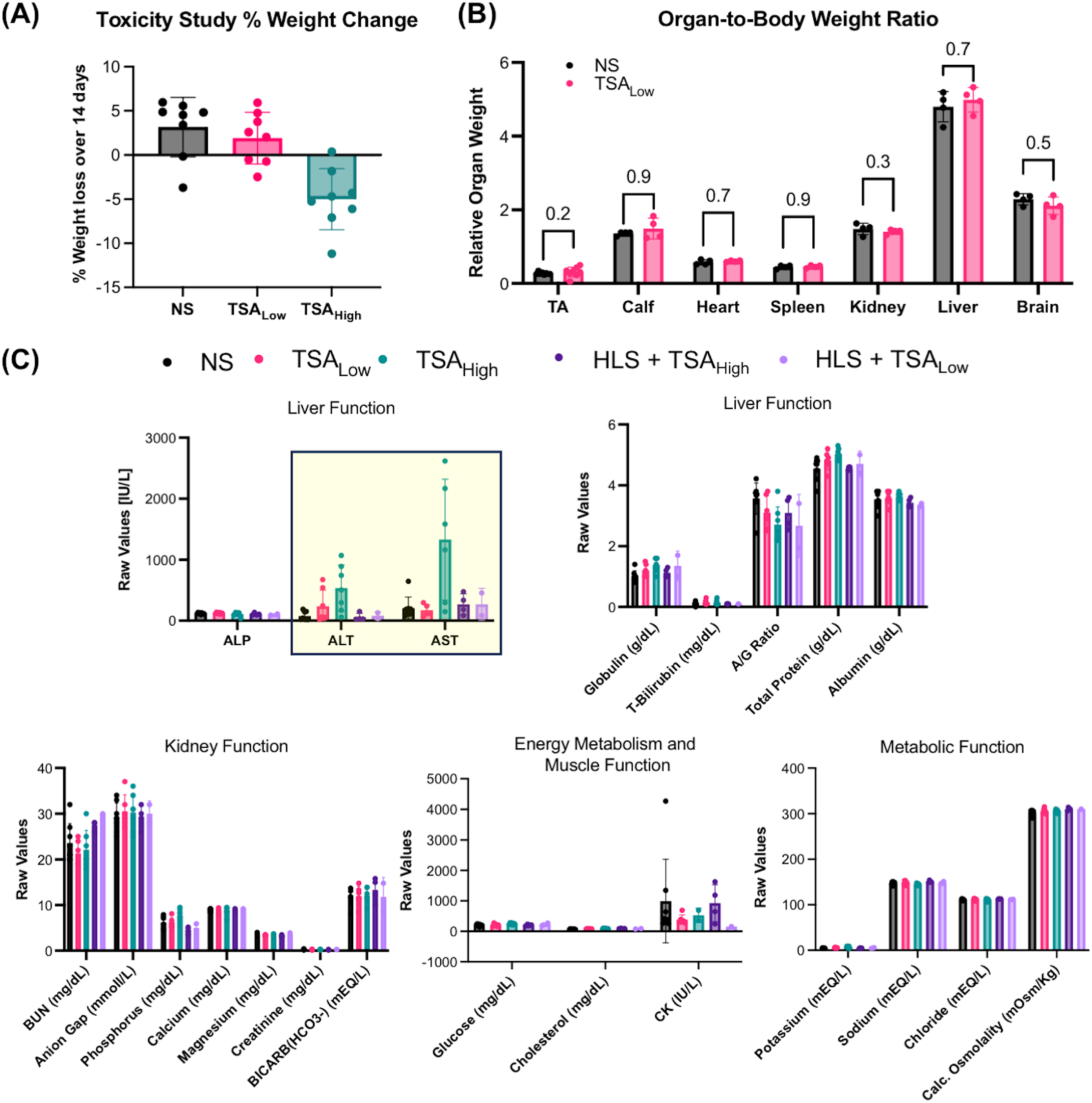
Assessment of systemic effects and potential toxicity following low- and high-dose Trichostatin A (TSA) administration. **(A)** Percentage change in body weight over 14 days in non-suspended mice (NS) receiving no treatment, NS + low-dose TSA (1.5 mg/kg), or NS + high-dose TSA (8 mg/kg). Low-dose TSA-treated mice exhibited body-weight changes comparable to untreated NS mice, whereas high-dose TSA-treated mice showed a trend toward body-weight loss. Error bars represent mean ± SD. **(B)** Organ-to-body weight ratios of the tibialis anterior (TA), calf muscle, heart, spleen, kidney, liver, and brain in NS mice with or without low-dose TSA treatment. All organs were dissected and individually weighed and then normalized to each rodents day 14 weight. No statistically significant differences were observed between groups. Error bars represent mean ± SD. **(C)** Serum biochemical analysis of liver function, kidney function, energy metabolism, muscle-associated enzymes, and electrolyte balance in NS mice receiving no treatment, low-dose TSA, or high-dose TSA, and in HLS + low TSA and HLS + high TSA mice. Most biochemical parameters remained broadly comparable across groups, although elevated alanine aminotransferase (ALT) and aspartate aminotransferase (AST) levels were observed in non-suspended mice receiving high-dose TSA. Error bars represent mean ± SD.

We first evaluated changes in body weight in non-suspended mice over the 14-day treatment period. Mice in the NS group exhibited a modest decrease in body weight, while mice receiving low-dose TSA showed a comparable trend. In contrast, non-suspended mice receiving high-dose TSA exhibited an average increase in body weight, suggesting that the higher drug concentration may have systemic effects independent of mechanical unloading (Fig. 2A). To further assess potential changes in tissue mass, we measured organ-to-body weight ratios in NS mice receiving low-dose TSA. No statistically significant differences were observed in the relative weights of the tibialis anterior, gastrocnemius, heart, spleen, kidney, liver, or brain compared with untreated NS mice (Fig. 2B), suggesting that low-dose TSA did not substantially alter the relative mass of these tissues over the treatment period.

We next performed a comprehensive serum biochemical analysis to assess liver and kidney function, metabolic activity, and electrolyte balance following TSA administration (Fig. 2C). Most measured parameters, including alkaline phosphatase (ALP), albumin, total protein, bilirubin, blood urea nitrogen (BUN), creatinine, glucose, and major electrolytes, remained broadly comparable across the experimental groups. However, non-suspended mice receiving high-dose TSA exhibited marked elevations in alanine aminotransferase (ALT) and aspartate aminotransferase (AST), with considerable variability among individual animals. These elevations were not similarly apparent in the HLS + high TSA group. Creatine kinase (CK) levels also showed substantial inter-animal variability, whereas the remaining metabolic and electrolyte measurements showed no prominent treatment-associated changes.

Collectively, these results suggest that low-dose TSA administration was relatively well tolerated over the 14-day experimental period, without substantial alterations in body weight, relative tissue mass, or the measured serum biochemical parameters. Although most serum markers remained comparable following high-dose TSA treatment, the elevated ALT and AST levels observed in non-suspended mice indicate a potential adverse effect that warrants further investigation. These findings highlight the importance of evaluating dose-dependent systemic effects of TSA. They also suggest that its therapeutic value lies in treating disuse-induced atrophy rather than enhancing performance, since liver enzymes rose in non-suspended TSA-treated mice but not in unloaded mice.

### TSA treatment partially preserves muscle architecture following hindlimb suspension

To investigate whether TSA-mediated preservation of locomotor function was accompanied by changes in muscle morphology, we examined histological sections of the femur and surrounding quadriceps muscle following 14 days of hindlimb suspension. Sagittal and transverse sections were stained with hematoxylin and eosin (H&E) and Masson’s trichrome to assess tissue architecture and connective tissue organization (Fig. 3).

**Figure 3.**
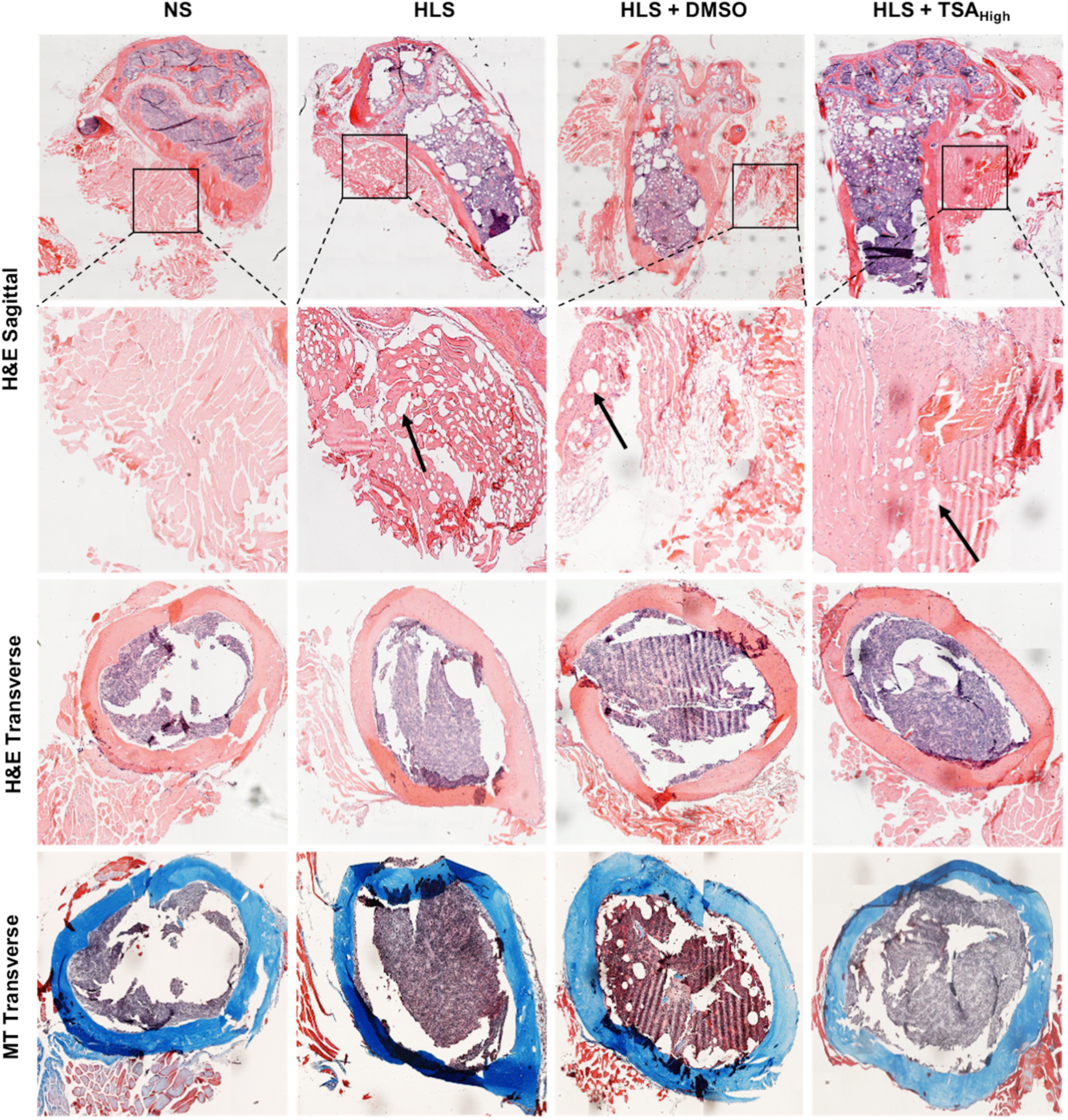
Histological assessment of quadriceps muscle and adjacent femoral bone following hindlimb suspension and TSA treatment. Representative histological images from NS, HLS, HLS + DMSO, and HLS + high TSA (8 mg/kg) mice after 14 days of hindlimb suspension. The first row shows sagittal H&E-stained sections of the distal femur and surrounding quadriceps muscle, with boxed regions indicating areas shown at higher magnification in the second row. HLS and HLS + DMSO mice exhibited disrupted muscle organization and prominent vacuole-like spaces (arrows), which appeared less pronounced following high-dose TSA treatment. The third and fourth rows show transverse sections of the femur stained with H&E and Masson’s trichrome, respectively, illustrating cortical bone and marrow architecture across the experimental groups.

H&E staining of sagittal sections revealed relatively intact and organized muscle fibers in the NS group, whereas HLS and HLS + DMSO mice exhibited noticeable disruption of muscle architecture. Higher-magnification images showed increased separation between muscle bundles and the presence of prominent vacuole-like spaces within the muscle tissue, suggesting unloading-associated tissue remodeling. In contrast, the HLS + high TSA group displayed more continuous muscle architecture, with fewer apparent vacuole-like spaces and better preservation of tissue organization. These observations suggest that HDAC inhibition partially attenuates the structural deterioration associated with prolonged mechanical unloading.

We further examined transverse sections of the femur using H&E and Masson’s trichrome staining. The gross cortical bone architecture appeared largely preserved across the experimental groups, without obvious structural disruption at the resolution examined. Masson’s trichrome staining revealed collagen-rich regions within the cortical bone, although no consistent treatment-associated differences could be established from these representative sections.

Overall, these histological observations suggest that high-dose TSA treatment partially preserves quadriceps muscle architecture following hindlimb suspension, consistent with the improved locomotor outcomes observed in TSA-treated mice (Fig. 1). These qualitative findings provide further evidence of tissue-level changes associated with TSA treatment and motivate a more detailed investigation of the underlying molecular mechanisms.

### HDAC inhibition shifts the skeletal muscle transcriptome toward the baseline state following hindlimb suspension

To investigate the molecular changes underlying the protective effects of TSA against disuse-induced muscle atrophy, we performed bulk RNA sequencing of calf muscles collected after 14 days of hindlimb suspension (n = 3 mice per group). We compared gene expression profiles among NS, HLS, HLS + DMSO, HLS + low TSA, and HLS + high TSA mice to determine the extent of transcriptional changes induced by mechanical unloading and their response to HDAC inhibition.

Principal component analysis (PCA) revealed distinct transcriptional profiles across the experimental groups (Fig. 4A). NS and HLS samples separated primarily along the first principal component (PC1), which accounted for 42% of the total variance, while PC2 accounted for an additional 27%. Samples from the HLS + DMSO and HLS + low TSA groups exhibited greater dispersion, suggesting heterogeneous transcriptional responses. In contrast, samples from the HLS + high TSA group occupied a more restricted region of the PCA space and shifted toward the NS group along PC1. Although the two groups remained separated along PC2, this distribution suggested that high-dose TSA partially restored the global transcriptional profile altered by mechanical unloading.

**Figure 4.**
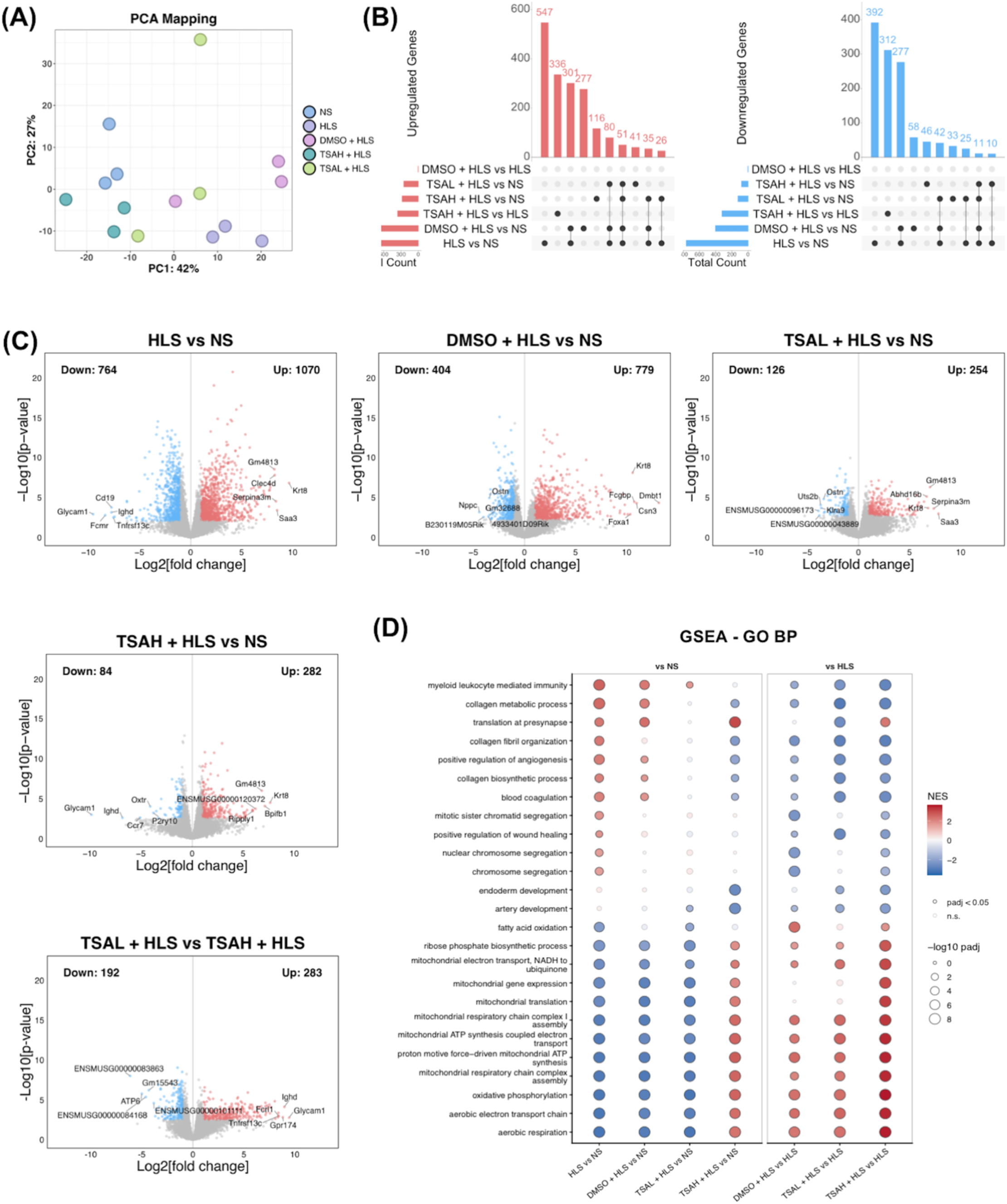
High-dose TSA treatment shifts the global skeletal muscle transcriptome toward the baseline state following hindlimb suspension. Bulk RNA sequencing was performed on calf muscle collected from NS, HLS, HLS + DMSO, HLS + low TSA (1.5 mg/kg), and HLS + high TSA (8 mg/kg) mice after 14 days of hindlimb suspension (n = 3 mice/group). **(A)** Principal component analysis (PCA) of global gene expression profiles. PC1 and PC2 account for 42% and 27% of the total variance, respectively. HLS samples separated from NS, while HLS + high TSA samples shifted toward the NS group along PC1. **(B)** UpSet plots showing the distribution and overlap of upregulated (left) and downregulated (right) genes across the indicated pairwise comparisons. **(C)** Volcano plots illustrating differential gene expression between HLS and NS, HLS + DMSO and NS, HLS + low TSA and NS, HLS + high TSA and NS, and HLS + low TSA and HLS + high TSA. Red and blue points indicate significantly upregulated and downregulated genes, respectively. The numbers of upregulated and downregulated genes are indicated in each plot. TSA-treated groups exhibited substantially fewer DEGs relative to NS than untreated HLS mice. **(D)** Gene set enrichment analysis (GSEA) of Gene Ontology Biological Process (GO BP) terms across the indicated comparisons relative to NS (left) and HLS (right). Dot color represents the normalized enrichment score (NES), with red indicating positive enrichment and blue indicating negative enrichment. Dot size corresponds to the −log10 adjusted *p*-value. High-dose TSA attenuated the enrichment of immune- and collagen-associated pathways while enhancing mitochondrial respiration, oxidative phosphorylation, and ATP synthesis-associated transcriptional programs.

Differential gene expression analysis further demonstrated the magnitude of transcriptional disruption following hindlimb suspension (Fig. 4B,C). Compared with NS, HLS muscle exhibited 1,834 differentially expressed genes (DEGs), including 1,070 upregulated and 764 downregulated genes. The HLS + DMSO group exhibited 1,183 DEGs relative to NS, comprising 779 upregulated and 404 downregulated genes. TSA treatment substantially reduced the number of genes differing from NS, with 380 DEGs (254 upregulated and 126 downregulated) in the HLS + low TSA group and 366 DEGs (282 upregulated and 84 downregulated) in the HLS + high TSA group. UpSet analysis revealed shared and comparison-specific DEGs across the treatment groups, while direct comparison of HLS + low TSA and HLS + high TSA identified 475 DEGs, indicating distinct transcriptional responses to the two TSA doses (Fig. 4B,C).

To determine whether the reduced transcriptional divergence reflected reversal of unloading-associated gene expression changes, we further examined the expression patterns of 3,161 DEGs identified across the experimental comparisons (Fig. S6). Heatmap analysis revealed distinct patterns of gene regulation among the experimental groups, with TSA treatment altering the direction and magnitude of numerous HLS-associated transcriptional changes. We then classified genes initially upregulated or downregulated by HLS according to the extent to which their expression changes were reversed following treatment. High-dose TSA produced the strongest overall restoration, with the majority of HLS-altered genes showing at least 50% reversal of their unloading-associated expression changes. Low-dose TSA exhibited an intermediate response, whereas a larger proportion of genes remained unchanged or only partially restored in the HLS + DMSO group (Fig. S6). These findings indicate that TSA treatment broadly counteracts the transcriptional response to mechanical unloading, with the more pronounced gene-level restoration observed at the higher dose.

We next performed GSEA usingGO BP terms to identify the biological programs associated with these transcriptional changes (Fig. 4D). Relative to NS, HLS muscle exhibited positive enrichment of pathways associated with immune responses, collagen metabolism, extracellular matrix organization, and tissue remodeling. In contrast, pathways related to mitochondrial electron transport, oxidative phosphorylation, ATP synthesis, and aerobic respiration were negatively enriched, consistent with suppression of mitochondrial metabolic programs following mechanical unloading. High-dose TSA substantially altered these enrichment patterns. Immune- and collagen-associated pathways were attenuated relative to HLS, while mitochondrial respiration and oxidative phosphorylation pathways became strongly positively enriched. Notably, several mitochondrial pathways also showed positive enrichment relative to NS, suggesting that TSA treatment may activate these metabolic programs beyond their baseline transcriptional state rather than simply restoring their original expression levels.

Analysis using the MSigDB Hallmark gene sets independently supported these findings (Fig. S6). HLS was associated with positive enrichment of inflammatory and stress-associated pathways, including TNFα signaling through NF-κB, IL6–JAK–STAT3 signaling, and epithelial–mesenchymal transition, together with negative enrichment of oxidative phosphorylation and fatty acid metabolism. High-dose TSA attenuated the inflammatory transcriptional signatures and strongly enriched oxidative phosphorylation relative to HLS, consistent with the GO BP analysis.

Together, these findings demonstrate that TSA treatment substantially modifies the global transcriptional response of skeletal muscle after mechanical unloading. The shift toward the NS transcriptomic profile, coupled with reversal of inflammatory and extracellular matrix-associated programs and enhancement of mitochondrial metabolic pathways, suggests that HDAC inhibition counteracts key molecular features of disuse-induced muscle deterioration. These transcriptional changes provide a potential molecular basis for the improved locomotor function and preservation of muscle architecture observed following TSA treatment.

### TSA attenuates FOXO-associated muscle atrophy programs and alters HDAC-related transcriptional responses

To identify the genes underlying the broad transcriptional changes observed following TSA treatment, we examined the expression of genes involved in muscle atrophy, protein degradation, and epigenetic regulation. Hierarchical clustering of selected genes revealed distinct expression profiles between NS and HLS muscle, with HLS + DMSO exhibiting a largely similar pattern to HLS (Fig. 5A). High-dose TSA shifted the expression of many of these genes toward the NS profile, whereas low-dose TSA produced a more intermediate response. Among the most prominent changes were genes associated with muscle protein degradation, stress responses, and myogenic regulation, suggesting that TSA treatment influences multiple components of the unloading-associated atrophy program.

**Figure 5.**
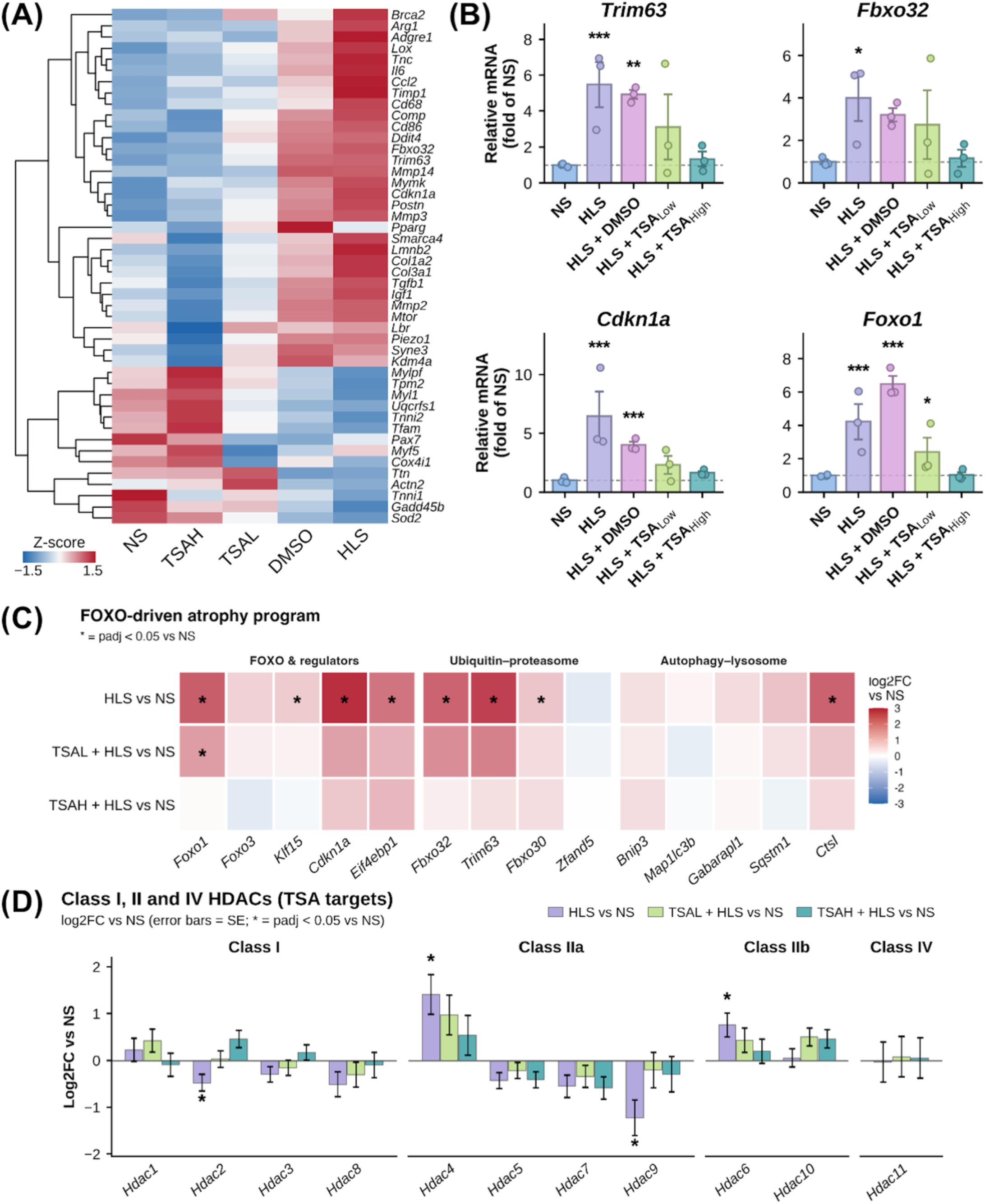
TSA treatment attenuates unloading-associated atrophy gene expression and alters HDAC-related transcriptional programs in skeletal muscle. **(A)** Hierarchically clustered heatmap of selected genes associated with muscle atrophy, inflammation, extracellular matrix remodeling, muscle structure, and transcriptional regulation across NS, HLS, HLS + DMSO, HLS + low TSA, and HLS + high TSA groups. Colors represent condition-averaged Z-scores, illustrating the reversal of multiple HLS-associated expression patterns following TSA treatment. **(B)** Relative transcript abundance of the atrophy-associated genes *Trim63, Fbxo32, Cdkn1a,* and *Foxo1*, expressed as fold change relative to NS. All four genes exhibited increased expression following HLS, with expression generally shifting toward NS levels following TSA treatment, particularly at the higher dose. **(C)** Heatmap showing log2 fold changes relative to NS for genes associated with FOXO regulation, ubiquitin – proteasome-mediated protein degradation, and autophagy-lysosome pathways. HLS was associated with increased expression of several atrophy-related genes, including *Foxo1, Cdkn1a, Trim63,* and *Fbxo32*, which was attenuated following TSA treatment. **(D)** RNA-seq-derived log2 fold changes relative to NS for histone deacetylase family members grouped into Class I (*Hdac1, Hdac2, Hdac3, Hdac8*), Class IIa (*Hdac4, Hdac5, Hdac7, Hdac9*), Class IIb (*Hdac6, Hdac10*), and Class IV (*Hdac11*). HLS induced differential transcriptional responses among HDAC family members, several of which shifted toward baseline expression following TSA treatment. RNA sequencing was performed using n = 3 mice/group. Error bars in quantitative plots represent standard error. Statistical annotations indicate comparisons with NS using DESeq2-adjusted *p*-values (Benjamini–Hochberg correction): * adjusted *p* < 0.05, ** adjusted *p* < 0.01, *** adjusted *p* < 0.001.

We first focused on the FOXO-associated transcriptional program, which regulates several genes involved in muscle protein degradation. HLS induced substantial increases in *Foxo1*, the muscle-specific E3 ubiquitin ligases *Trim63* (MuRF1) and *Fbxo32* (Atrogin-1), and the cell cycle regulator *Cdkn1a* (p21) compared with NS (Fig. 5B). These changes were largely retained in HLS + DMSO mice but progressively attenuated following TSA treatment. Notably, high-dose TSA reduced the expression of all four genes toward baseline levels, while low-dose TSA produced an intermediate response. The increase in *Foxo1* expression was particularly pronounced in HLS + DMSO muscle, whereas high-dose TSA restored its transcript abundance close to that of NS. These findings suggest that TSA treatment counteracts the activation of a major transcriptional program associated with disuse-induced muscle wasting.

We next examined genes involved in the ubiquitin-proteasome and autophagy-lysosome pathways to determine whether the response extended beyond the canonical atrophy markers. HLS was associated with increased expression of several FOXO-related and proteolytic genes, including *Klf15*, *Eif4ebp1*, *Fbxo30*, and *Ctsl*, in addition to *Trim63* and *Fbxo32* (Fig. 5C). The expression of these genes generally shifted toward NS levels following TSA treatment, with a more pronounced response at the higher dose. However, not all genes associated with autophagy exhibited substantial changes, indicating that unloading and TSA treatment did not uniformly affect every component of the protein degradation machinery. The broader panel of atrophy-associated genes showed a similar pattern, with HLS-induced increases in *Runx1*, *Ddit4*, *Gadd45a*, and several other stress-associated transcripts attenuated following TSA treatment (Fig. S7A). Additional quantitative analysis of FOXO-associated genes independently supported these trends (Fig. S12A).

Because TSA inhibits HDAC activity, we next investigated whether mechanical unloading and TSA treatment were accompanied by altered transcription of HDAC family members. HLS induced distinct responses among individual HDACs, including increased expression of *Hdac4* and *Hdac6* and reduced expression of *Hdac2* and *Hdac9* relative to NS (Fig. 5D). TSA treatment modified several of these expression patterns, with *Hdac4*, *Hdac6*, and *Hdac9* generally shifting toward baseline levels following high-dose treatment. Additional profiling of chromatin-associated regulatory genes revealed unloading- and treatment-associated changes in several histone modifiers, chromatin remodeling factors, and epigenetic regulators (Fig. S9). These included changes in the expression of *Tet1*, *Smarcd3*, and several other genes involved in chromatin organization. The broader profiles of Class I, II, and IV HDAC family members are presented in Fig. S12C.

We further examined genes associated with the HDAC4-myogenic regulatory network, including members of the MEF2 family and genes involved in muscle differentiation and remodeling. HLS increased the expression of several injury- and remodeling-associated genes, particularly *Myog*, *Chrna1*, *Runx1*, and *Gadd45a*, while the expression of individual MEF2 family members showed comparatively modest changes (Fig. S12B,D). High-dose TSA attenuated the expression of several of these HLS-responsive genes, consistent with a broader shift in myogenic and stress-associated transcriptional regulation.

Together, these results demonstrate that mechanical unloading induces a coordinated transcriptional response involving FOXO-associated atrophy genes, protein degradation machinery, and epigenetic regulators. TSA treatment, particularly at the higher dose, substantially attenuates several components of this response. Although changes in HDAC transcript abundance do not directly establish altered HDAC enzymatic activity, the transcriptional findings are consistent with the involvement of HDAC-sensitive regulatory pathways in the muscle response to mechanical unloading.

### TSA broadly remodels inflammatory, extracellular matrix, contractile, and metabolic gene programs following mechanical unloading

The global enrichment analysis in Figure 4 indicated that the transcriptional consequences of HLS extended beyond muscle protein degradation to include inflammatory signaling, extracellular matrix remodeling, and mitochondrial metabolism. To examine these responses in greater detail, we analyzed selected genes associated with these biological processes and compared their expression across the experimental groups.

Mechanical unloading induced a pronounced inflammatory transcriptional signature characterized by increased expression of genes associated with immune cell recruitment, macrophage-associated responses, and inflammatory signaling. These included *Il6*, *Ccl2*, *Ccl7*, *Cd68*, *Itgam*, *Trem2*, *S100a8*, and *S100a9*, among others (Fig. S7B). Many of these genes remained elevated in HLS + DMSO muscle, whereas their expression was generally attenuated following TSA treatment. The reduction was particularly apparent in the HLS + high TSA group, although some inflammatory transcripts remained elevated relative to NS. These findings support the GSEA results showing attenuation of immune-associated transcriptional programs following HDAC inhibition (Fig. 4D and Fig. S6B).

A similar pattern was observed among genes associated with extracellular matrix organization and tissue remodeling. HLS increased the expression of several matrix-associated genes, including *Tnc*, *Timp1*, *Postn*, *Lox*, *Mmp3*, *Thbs1*, and *Col1a1*, suggesting an enhanced transcriptional response involving matrix deposition, turnover, and tissue remodeling (Fig. S10A). These changes were partially retained in HLS + DMSO muscle but were generally reduced following TSA treatment. We also examined genes associated with fibro-adipogenic progenitor-related processes, including *Postn*, *S100a4*, *Ccn4*, and several collagen-associated genes, which showed similar unloading-associated increases and treatment-dependent attenuation (Fig. S8B). These patterns suggest that TSA treatment counteracts transcriptional programs associated with inflammatory and extracellular matrix remodeling during muscle disuse.

We next evaluated genes involved in muscle structure, contractile function, and myogenic differentiation. HLS was associated with increased expression of several developmental and remodeling-associated muscle genes, including *Myh3*, *Myh8*, and *Tnnt2*, accompanied by reduced expression of several genes associated with mature muscle contractile architecture, including *Acta1*, *Tnnt1*, *Ldb3*, *Tpm1*, and *Myl1* (Fig. S7C). TSA treatment, particularly at the higher dose, attenuated many of these changes. Additional muscle-associated genes, including *Myog*, *Myf6*, and *Chrna1*, showed unloading-associated transcriptional responses that were also modified by TSA treatment (Fig. S7D). Analysis of genes associated with satellite cell differentiation and myogenic regulation revealed increased expression of *Mymk*, *Mymx*, and *Cdkn1a* following HLS, with several of these changes reduced after TSA treatment (Fig. S8C). These findings suggest that the transcriptional response to unloading involves altered regulation of muscle structural maintenance and myogenic remodeling, both of which are responsive to HDAC inhibition.

Genes associated with anabolic signaling and metabolic regulation were also investigated. HLS increased the expression of several genes involved in the regulation of muscle growth and stress responses, including *Ddit4*, *Eif4ebp1*, and *Pik3r1* (Fig. S8A). Notably, *Ddit4* and *Eif4ebp1* are associated with the suppression of mTOR-dependent protein synthesis, suggesting that their increased expression may contribute to the altered anabolic environment of unloaded muscle. These transcriptional changes were generally attenuated following TSA treatment, although the expression of individual signaling components varied across groups.

Consistent with the pathway-level analysis, HLS also altered the expression of genes associated with mitochondrial metabolism and cellular energy regulation. Several stress- and metabolic adaptation-associated genes, including *Hmox1*, *Foxo1*, and *Pdk4*, were upregulated, whereas genes involved in mitochondrial respiratory function, including *Ndufa9*, *Sdha*, *Pdha1*, and *Cox4i1*, showed reduced expression relative to NS (Fig. S10B). TSA treatment shifted the expression of several of these genes toward baseline levels, particularly at the higher dose. These findings complement the strong positive enrichment of oxidative phosphorylation and mitochondrial respiration pathways observed following high-dose TSA treatment (Fig. 4D), suggesting a broad reorganization of metabolic gene expression during the response to HDAC inhibition.

Finally, given the mechanical origin of disuse atrophy, we examined the transcriptional response of genes associated with mechanotransduction and nuclear architecture. HLS increased the expression of several mechanically responsive genes, including *Ankrd1*, *Ccn1*, *Ccn2*, and *Itga7*, with varying degrees of attenuation following TSA treatment (Fig. S11A). Genes associated with the nuclear envelope and nuclear–cytoskeletal coupling, including *Syne1*, *Lmna*, and *Sun2*, also exhibited unloading-associated transcriptional changes, although their responses to TSA were not uniform (Fig. S11B). These observations suggest that mechanical unloading affects not only conventional muscle metabolic and contractile pathways but also transcriptional programs associated with cellular force sensing and nuclear organization.

Collectively, these gene-level analyses demonstrate that TSA treatment modifies a broad range of biological processes affected by mechanical unloading. TSA treatment dampened inflammatory, extracellular matrix, and atrophy-related gene programs while restoring structural and mitochondrial genes, providing a molecular basis for the functional and histological improvements observed.

### Dose-dependent reversal of unloading-associated gene expression reveals distinct molecular programs of TSA-mediated rescue

Although both TSA doses reduced the overall transcriptional divergence from NS (Fig. 4), the preceding analyses suggested that individual genes and biological pathways differed in their responsiveness to treatment. We therefore performed a quantitative analysis of transcriptional rescue to determine the extent to which low- and high-dose TSA reversed the changes induced by mechanical unloading and to identify distinct patterns of gene regulation.

We first examined the 1,834 genes differentially expressed between HLS and NS and compared their HLS-induced log2 fold changes with the corresponding expression changes in HLS + DMSO, HLS + low TSA, and HLS + high TSA (Fig. 6A). In this analysis, a regression slope approaching one indicates persistence of the unloading-associated transcriptional response, whereas a slope approaching zero indicates greater restoration toward NS. The HLS + DMSO group exhibited a slope of 0.72, corresponding to an estimated 28% global reversal of the HLS-associated expression changes. The slope decreased to 0.49 following low-dose TSA and further to 0.39 following high-dose TSA, corresponding to estimated global reversals of 51% and 61%, respectively. These findings indicate that both TSA doses counteracted the transcriptional response to unloading, with the larger overall effect observed following high-dose treatment.

**Figure 6.**
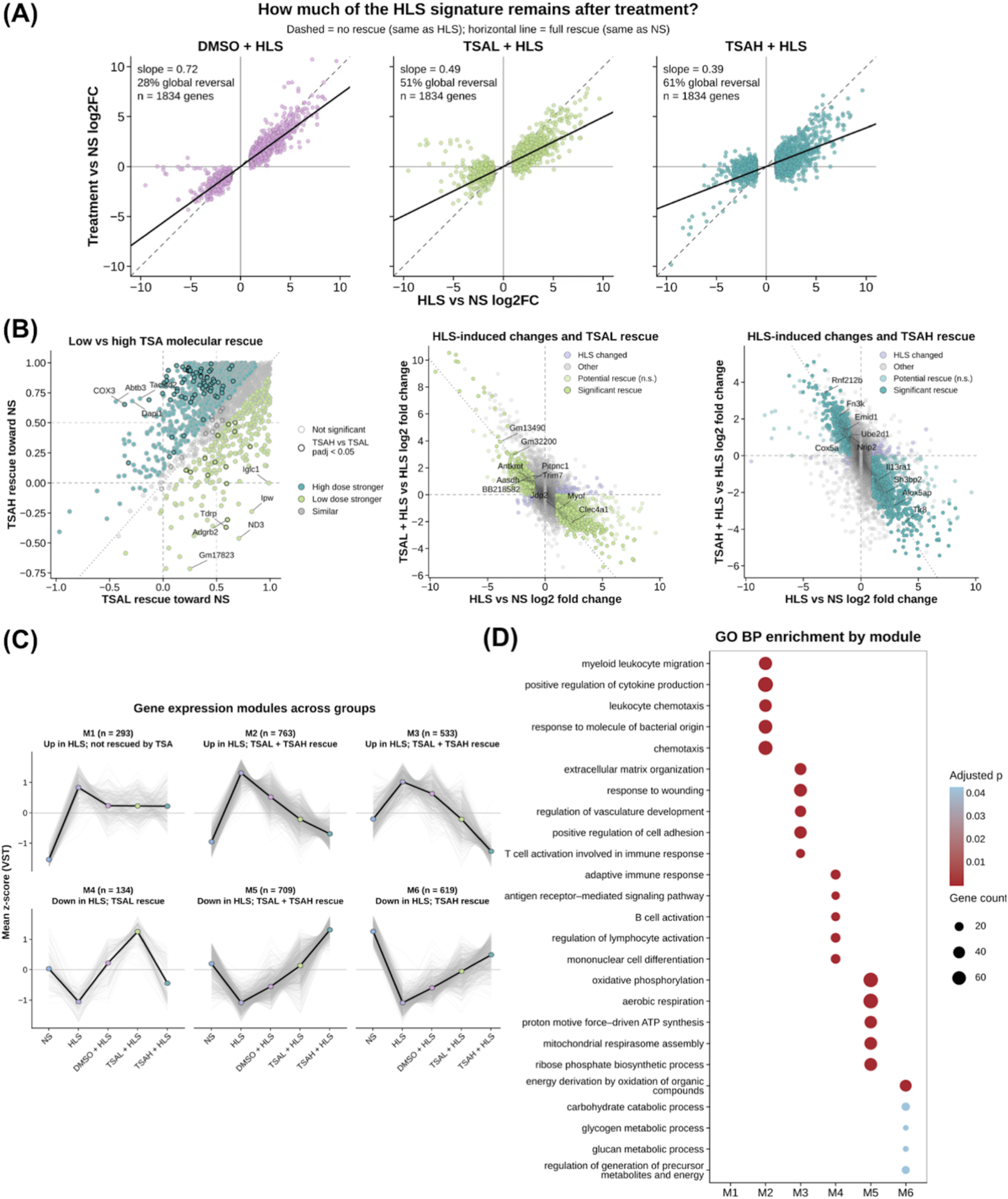
Dose-dependent reversal of unloading-associated transcriptional changes reveals distinct molecular programs responsive to TSA treatment. **(A)** Genome-wide comparison of HLS-induced transcriptional changes and their response to treatment. For 1,834 genes differentially expressed between HLS and NS, log2 fold changes following HLS are plotted against the corresponding changes in HLS + DMSO, HLS + low TSA, and HLS + high TSA groups relative to NS. Dashed diagonal lines indicate persistence of the HLS-associated expression changes, whereas values approaching zero on the vertical axis indicate restoration toward NS. Linear regression slopes of 0.72, 0.49, and 0.39 correspond to estimated global reversal of 28%, 51%, and 61% following DMSO, low-dose TSA, and high-dose TSA, respectively, calculated as 1 − slope. **(B)** Gene-level comparison of transcriptional rescue following low- and high-dose TSA. Left: scatterplot comparing the degree of rescue toward NS between TSA doses, with genes classified according to the dose producing the stronger response. Outlined points indicate significant differences between doses (adjusted *p* < 0.05). Middle and right: scatterplots showing HLS-induced log2 fold changes relative to NS against treatment-induced changes relative to HLS for low- and high-dose TSA, respectively. Genes are classified according to whether they exhibit significant or potential transcriptional rescue. **(C)** Unbiased grouping of genes using k-means clustering into six expression modules based on their responses to HLS and TSA treatment. Modules represent genes upregulated by HLS that remain incompletely restored (M1, n = 293), upregulated genes responsive to both TSA doses (M2, n = 763; M3, n = 533), downregulated genes preferentially responsive to low-dose TSA (M4, n = 134), downregulated genes responsive to both doses (M5, n = 709), and downregulated genes preferentially responsive to high-dose TSA (M6, n = 619). Thin lines represent individual gene-expression profiles, and bold lines indicate module-average expression patterns. **(D)** GO BP enrichment analysis of the six expression modules. Dot size represents the number of genes associated with each enriched biological process, while color indicates the adjusted *p*-value. The modules distinguish immune and inflammatory responses, extracellular matrix remodeling, mitochondrial oxidative phosphorylation, and carbohydrate metabolism-associated transcriptional programs.

We next asked whether the greater global reversal with high-dose TSA reflected uniform improvement across genes or selective effects on particular targets. To address this, we compared the degree of recovery toward NS for each gene following low- and high-dose treatment (Fig. 6B, left). Although many genes recovered to a similar extent with both doses, distinct subsets responded more strongly to one dose than the other, and 102 genes differed significantly between the two treatment groups. Thus, increasing the TSA dose did not simply amplify the low-dose response. We then plotted each gene’s HLS-induced change against its change following treatment relative to HLS (Fig. 6B, middle and right). With both doses, many genes shifted in the opposite direction from their unloading-induced change, consistent with reversal toward NS, although the magnitude and statistical significance of these shifts varied between genes. Together, these analyses indicate that TSA-mediated transcriptional rescue is heterogeneous across genes rather than a uniform reduction of the unloading response.

To identify broader patterns underlying this heterogeneity, we performed data-driven k-means clustering of the responsive genes based on their expression across NS, HLS, HLS + DMSO, HLS + low TSA, and HLS + high TSA groups. This analysis identified six distinct expression modules (M1-M6), representing genes with different responses to unloading and TSA treatment (Fig. 6C).

Three modules contained genes that were predominantly upregulated following HLS. Module M1 (n = 293) consisted of genes whose expression remained elevated after TSA treatment, indicating a transcriptional response that was not fully reversed by either dose. In contrast, modules M2 (n = 763) and M3 (n = 533) contained HLS-induced genes whose expression was substantially reduced following TSA treatment. Although both doses affected these modules, the magnitude and pattern of recovery differed, with high-dose TSA producing a more pronounced reduction in several gene clusters.

The remaining three modules contained genes that were predominantly downregulated following HLS. Module M4 (n = 134) showed a preferential response to low-dose TSA, including a pronounced increase in expression beyond the NS-associated level in the module-average profile. Module M5 (n = 709) exhibited restoration with both TSA doses, with a particularly strong increase following high-dose treatment. Module M6 (n = 619) showed a more progressive dose-associated recovery, with high-dose TSA producing the greatest shift toward or beyond the NS expression level. These distinct profiles further demonstrate that the transcriptional consequences of HDAC inhibition vary across gene populations, with some programs responding to both doses and others exhibiting greater sensitivity to one treatment condition.

We next performed Gene Ontology Biological Process enrichment analysis to determine whether these expression modules represented distinct biological functions (Fig. 6D). Module M2 was strongly enriched in immune and inflammatory processes, including myeloid leukocyte migration, cytokine production, and leukocyte chemotaxis. Module M3 was enriched in extracellular matrix organization, wound responses, cell adhesion, and tissue remodeling-associated processes. Module M4 was associated predominantly with adaptive immune responses, including lymphocyte activation and antigen receptor-mediated signaling. In contrast, module M5 was strongly enriched in mitochondrial metabolic processes, including oxidative phosphorylation, aerobic respiration, mitochondrial respirasome assembly, and ATP synthesis. Module M6 contained genes associated with energy metabolism, carbohydrate catabolism, and glycogen and glucan metabolic processes.

The functional identities of these modules provide further insight into the dose-dependent response to TSA. The attenuation of HLS-induced immune and extracellular matrix-associated transcriptional programs was evident across treatment-responsive modules, whereas restoration of mitochondrial and energy metabolism-associated genes showed particularly prominent responses to high-dose TSA. These findings are consistent with the pathway-level enrichment analyses in Figure 4 and the individual gene-expression patterns described above.

Together, these analyses demonstrate that TSA treatment produces a substantial but selective reversal of the skeletal muscle transcriptional response to mechanical unloading. High-dose TSA provides the greatest overall reduction in unloading-associated expression changes, particularly among genes involved in mitochondrial metabolism and energy regulation, while other transcriptional programs exhibit comparable or preferential responses to low-dose treatment. Thus, HDAC inhibition does not simply return the muscle transcriptome uniformly to its baseline state but produces a dose-dependent reorganization of specific molecular programs associated with muscle atrophy, tissue remodeling, and metabolic adaptation.

### HDAC inhibition partially restores nuclear architecture and heterochromatin organization following mechanical unloading

Unloading altered the expression of numerous epigenetic regulators and mechanotransduction-associated genes, including components of the nuclear envelope and the LINC complex that physically links the cytoskeleton to the nucleus (Figs. 5D, S9, and S11). Because these structures help convert mechanical cues into changes in chromatin organization, we asked whether the transcriptional changes were accompanied by structural changes in the nucleus. The nuclear lamina, a meshwork of lamin proteins beneath the inner nuclear membrane, maintains nuclear shape and anchors transcriptionally silent heterochromatin at the nuclear periphery. Much of this peripheral heterochromatin is marked by histone H3 lysine 9 trimethylation (H3K9me3). Changes in lamina organization can therefore alter both nuclear morphology and the distribution of H3K9me3-marked chromatin. We therefore used super-resolution immunofluorescence microscopy of calf muscle sections to examine Lamin A/C organization, as a readout of nuclear lamina structure, and H3K9me3, as a readout of heterochromatin organization. In NS muscle, nuclei exhibited an elongated morphology with a relatively continuous Lamin A/C distribution along the nuclear periphery (Fig. 7A). In contrast, nuclei from HLS mice displayed heterogeneous morphological abnormalities, including irregular nuclear contours, apparent constrictions, and altered Lamin A/C organization. These changes were particularly evident in nuclei exhibiting pronounced deformation and irregular folding of the nuclear envelope. Following high-dose TSA treatment, the representative nuclei exhibited a more continuous peripheral Lamin A/C distribution and a comparatively regular morphology, suggesting partial preservation of nuclear architecture despite continued mechanical unloading.

**Figure 7.**
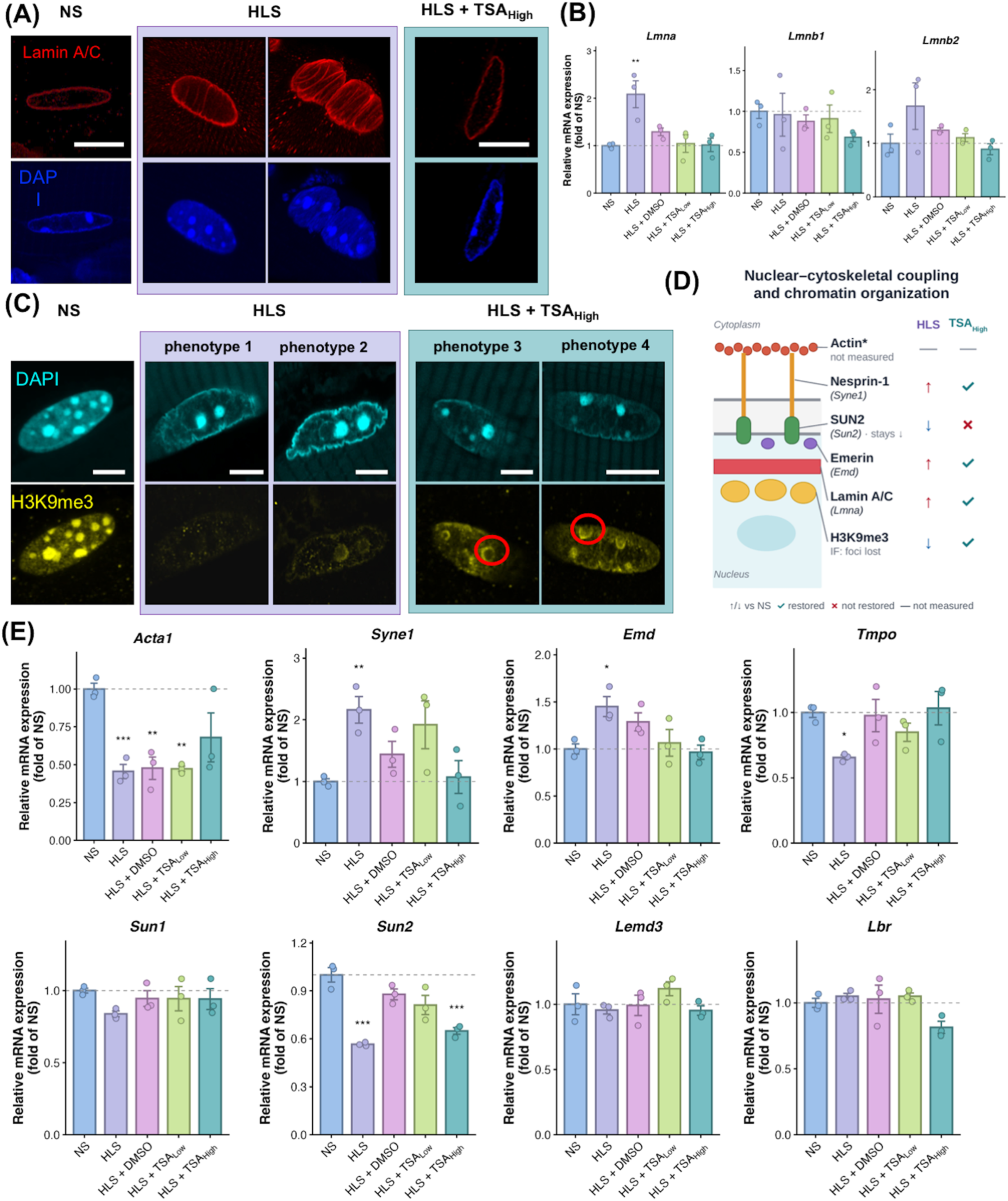
HDAC inhibition partially restores nuclear lamina architecture and H3K9me3-associated heterochromatin organization following hindlimb suspension. **(A)** Representative super-resolution immunofluorescence images of nuclei in calf muscle sections stained for Lamin A/C (red) and DAPI (blue). NS nuclei exhibited relatively continuous peripheral Lamin A/C organization, whereas HLS nuclei showed irregular nuclear morphology, apparent constrictions, and altered Lamin A/C distribution. High-dose TSA treatment partially preserved the continuity of the nuclear lamina. Scale bars: 5 µm. **(B)** Relative mRNA expression of nuclear lamina-associated genes *Lmna*, *Lmnb1*, and *Lmnb2* across NS, HLS, HLS + DMSO, HLS + low TSA, and HLS + high TSA groups. *Lmna* expression was significantly elevated following HLS and shifted toward NS levels following TSA treatment. **(C)** Representative immunofluorescence images showing DAPI (cyan) and H3K9me3 (yellow) distribution in nuclei from NS, HLS, and HLS + high TSA mice. NS nuclei exhibited prominent H3K9me3-enriched foci, whereas HLS nuclei displayed reduced focal organization and more dispersed staining patterns (phenotypes 1 and 2). High-dose TSA treatment was associated with partial restoration of discrete H3K9me3-enriched regions (phenotypes 3 and 4; red circles). **(D)** Schematic illustrating selected components associated with nuclear–cytoskeletal coupling and nuclear envelope organization, including actin, Nesprin-1, SUN2, emerin, Lamin A/C, and H3K9me3-associated heterochromatin. Arrows indicate changes following HLS relative to NS, while symbols summarize the extent to which the observed transcriptional or imaging phenotype shifts toward baseline following high-dose TSA treatment. **(E)** Relative mRNA expression of selected cytoskeletal and nuclear envelope-associated genes, including *Acta1, Syne1, Emd, Tmpo, Sun1, Sun2, Lemd3,* and *Lbr*, across the five experimental groups. HLS altered the expression of several nuclear structural genes, with partial normalization following TSA treatment. Notably, *Sun2* expression remained reduced following high-dose TSA treatment. For quantitative gene-expression analyses, values are expressed relative to NS. Asterisks indicate significant differences from NS, as annotated in the plots.

Interestingly, these structural changes were accompanied by altered expression of nuclear lamina-associated genes. *Lmna* expression increased approximately twofold in HLS muscle relative to NS, whereas expression in the TSA-treated groups was closer to baseline (Fig. 7B). *Lmnb2* exhibited a similar, although less pronounced, increase following HLS, while *Lmnb1* expression remained comparatively stable across groups. The increased *Lmna* transcript abundance in unloaded muscle, despite the apparent disruption of Lamin A/C organization, suggests that transcriptional regulation of nuclear lamina components does not necessarily correspond to preservation of their structural organization. One possibility is that elevated *Lmna* expression reflects an adaptive response to altered nuclear mechanical loading, although this remains to be established experimentally.

We next examined the spatial distribution of H3K9me3, a histone modification associated with constitutive heterochromatin. In NS muscle, H3K9me3 staining was organized into prominent, discrete nuclear foci (Fig. 7C). Following HLS, this organization was markedly altered, with reduced prominence of discrete H3K9me3 foci and a more dispersed staining pattern. Representative HLS nuclei exhibited different degrees of this disruption, ranging from a substantial loss of distinguishable foci to a redistribution of H3K9me3-associated signal within the nucleus. In contrast, nuclei from HLS + high TSA mice exhibited a partial reappearance of discrete H3K9me3-enriched regions, although their distribution remained different from that observed in NS muscle. These findings suggest that HDAC inhibition mitigates unloading-associated alterations in heterochromatin organization, potentially contributing to the broader transcriptional changes observed following TSA treatment.

To further investigate the nuclear structural response to unloading, we examined the expression of genes associated with nuclear-cytoskeletal coupling, nuclear envelope organization, and structural maintenance (Fig. 7D,E). HLS significantly increased the expression of *Syne1*, encoding Nesprin-1, and *Emd*, encoding emerin, while reducing the expression of *Acta1*, *Tmpo*, and *Sun2*. High-dose TSA shifted the expression of *Syne1*, *Emd*, and *Tmpo* toward NS levels and partially attenuated the reduction in *Acta1*. However, *Sun2* expression remained significantly reduced following high-dose TSA treatment, indicating that the transcriptional response of nuclear envelope components was not uniformly restored. Other nuclear-associated genes, including *Sun1*, *Lemd3*, and *Lbr*, exhibited comparatively modest changes across the experimental groups. These observations are consistent with the broader transcriptomic analysis of mechanotransduction and nuclear envelope-associated genes (Fig. S11), suggesting that mechanical unloading affects multiple components involved in maintaining nuclear structure and its coupling to the cytoskeleton.

Together, these findings show that mechanical unloading alters nuclear morphology, Lamin A/C organization, and the distribution of H3K9me3-marked heterochromatin in skeletal muscle. High-dose TSA partially reversed these structural changes and returned the expression of several nuclear envelope genes toward NS levels, although *Sun2* remained reduced. Combined with the transcriptional changes described above, these results link HDAC-sensitive regulation to nuclear architecture remodeling in the skeletal muscle response to unloading.

#### TSA alters muscle-associated bone signaling programs without preventing early trabecular bone loss following mechanical unloading

Mechanical unloading causes loss of both skeletal muscle and bone, and the two tissues are closely linked through mechanical loading and secreted signaling factors. We therefore asked whether the muscle-protective effects of TSA extended to bone. After 14 days of hindlimb suspension, we assessed femoral architecture by micro-computed tomography (µCT) and bone mechanical properties by three-point bending in NS, HLS, and HLS + high-dose TSA mice (Fig. 8A-C).

**Figure 8.**
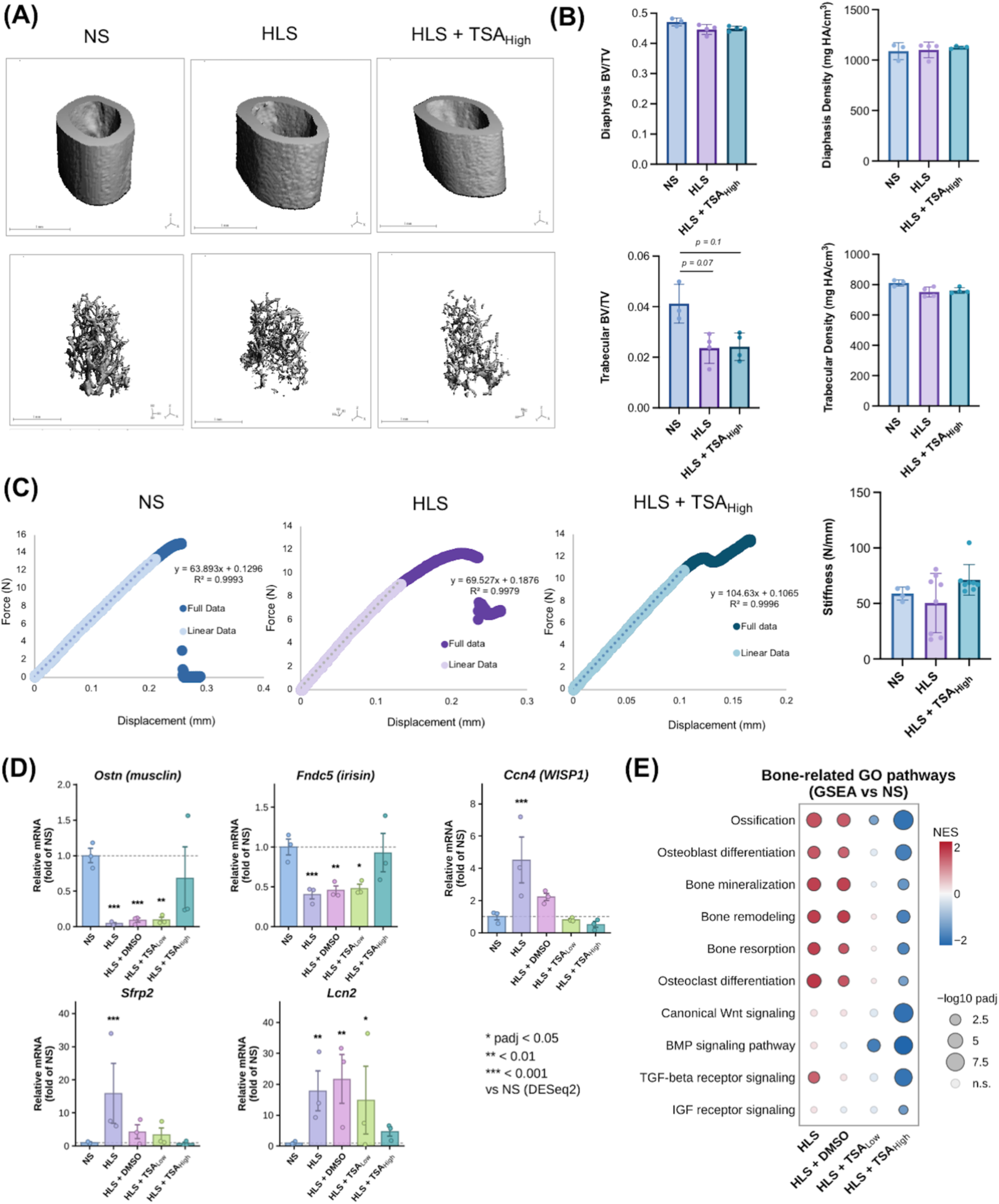
TSA modifies muscle-associated bone signaling programs without preventing early trabecular bone loss following hindlimb suspension. **(A)** Representative three-dimensional µCT reconstructions of the femoral cortical diaphysis (top) and trabecular bone compartment (bottom) from NS, HLS, and HLS + high TSA mice after 14 days of hindlimb suspension. **(B)** Quantification of diaphyseal bone volume fraction (BV/TV) and bone mineral density, together with trabecular BV/TV and bone mineral density. Diaphyseal measurements were comparable across groups, whereas trabecular BV/TV was reduced in HLS (*p* = 0.07) and HLS + high TSA (*p* = 0.1) compared with NS. No significant difference in trabecular BV/TV was detected between HLS and HLS + high TSA. Error bars represent mean ± SD and groups were compared by Kruskal–Wallis test followed by Dunn’s multiple comparisons test. **(C)** Representative force-displacement curves from femoral three-point bending tests, showing the linear regions of the ramp to failure portions used to calculate stiffness. Quantification of whole-femur bending stiffness revealed no statistically significant differences among groups. **(D)** Relative mRNA expression of selected genes associated with muscle-derived signaling and bone regulation, including *Ostn* (musclin), *Fndc5* (irisin precursor), *Ccn4* (WISP1), *Sfrp2*, and *Lcn2*. Expression values were obtained from calf muscle RNA sequencing and are shown relative to NS across the five experimental groups. Asterisks indicate DESeq2-adjusted significance relative to NS: * adjusted *p* < 0.05, ** adjusted *p* < 0.01, *** adjusted *p* < 0.001. Error bars represent standard error. **(E)** GSEA of bone-related GO BP terms using calf muscle RNA-seq data. Enrichment patterns are shown for HLS, HLS + DMSO, HLS + low TSA, and HLS + high TSA relative to NS. Dot color indicates the normalized enrichment score (NES), with red and blue representing positive and negative enrichment, respectively. Dot size corresponds to the −log10 adjusted *p*-value.

Quantitative µCT analysis revealed distinct responses in the cortical and trabecular compartments (Fig. 8B). No significant differences were detected in diaphyseal bone volume fraction (BV/TV) or mineral density among the experimental groups. In contrast, trabecular BV/TV was reduced in HLS mice compared with NS (*p* = 0.07), indicating that trabecular bone was particularly sensitive to the early effects of mechanical unloading. High-dose TSA treatment did not prevent this deterioration, as trabecular BV/TV remained lower than NS (*p* = 0.1) and was not significantly different from untreated HLS. Trabecular bone mineral density also remained comparable across groups. These findings demonstrate that the structural consequences of unloading were more pronounced in the trabecular compartment and that systemic TSA treatment did not measurably preserve trabecular bone volume over the 14-day experimental period.

We next evaluated whether these architectural changes affected the mechanical properties of the femur using three-point bending tests (Fig. 8C). Representative force-displacement curves showed an initial linear elastic response followed by nonlinear deformation, allowing stiffness to be determined from the linear region. Although the mean stiffness was lower in HLS mice and higher in HLS + high TSA mice, the differences were not statistically significant. The absence of a significant change in whole-femur bending stiffness, despite the reduction in trabecular BV/TV, may reflect the relatively preserved cortical architecture during this early period of unloading, as cortical geometry contributes substantially to whole-bone bending resistance.

Because skeletal muscle and bone communicate through mechanical loading and biochemical signaling, we next investigated whether TSA altered the expression of muscle-derived factors implicated in bone regulation. Using the calf muscle RNA-seq dataset, we examined several genes encoding secreted proteins or signaling regulators associated with muscle-bone interactions (Fig. 8D). HLS markedly reduced the expression of *Ostn*, which encodes musclin, and *Fndc5*, which encodes the precursor of irisin. Both genes have been implicated in musculoskeletal signaling and adaptation. Their expression remained suppressed in HLS + DMSO muscle but shifted toward NS levels following high-dose TSA treatment, although the response of *Ostn* exhibited considerable variability. In contrast, HLS induced substantial increases in *Ccn4* (WISP1), *Sfrp2*, and *Lcn2*, which encode proteins associated with extracellular matrix signaling, Wnt pathway regulation, and inflammatory or metabolic responses. These increases were attenuated following high-dose TSA treatment, with the expression of *Ccn4* and *Sfrp2* approaching baseline levels. Together, these findings show that unloading markedly alters muscle expression of factors that can signal to bone, and that HDAC inhibition reverses several of these changes.

To examine these responses more broadly, we analyzed genes involved in muscle-derived signaling, bone-associated receptors, Wnt/β-catenin regulation, TGF-β/BMP signaling, and pathways associated with osteogenic and inflammatory responses (Fig. S13). HLS induced substantial changes across these gene families, including reduced expression of several putative muscle-derived signaling factors and altered expression of Wnt pathway regulators. TSA treatment modified many of these transcriptional responses, although the effects were heterogeneous across individual genes and signaling pathways. Gene set enrichment analysis further revealed significant changes in biological processes annotated to ossification, osteoblast differentiation, bone remodeling, bone resorption, and Wnt and BMP signaling (Fig. 8E). In HLS muscle, several bone-related gene sets were positively enriched relative to NS, whereas high-dose TSA produced negative enrichment of multiple corresponding gene sets. These changes indicate substantial remodeling of bone-associated transcriptional programs within skeletal muscle, rather than directly demonstrating changes in bone cell activity.

Overall, 14 days of unloading reduced trabecular bone volume without detectable changes in diaphyseal architecture or whole-femur bending stiffness. Interestingly, although high-dose TSA substantially altered the expression of muscle-derived signaling factors and pathways linked to muscle–bone communication, it did not prevent this early trabecular loss. Several factors may explain this. First, bone remodeling is slower than the muscle transcriptional response, so a 14-day unloading period may be too short to detect downstream skeletal effects. Second, because TSA was given systemically, these experiments cannot separate indirect effects acting through muscle from direct effects of the drug on bone cells. These results therefore suggest that HDAC inhibition may modify signaling between muscle and bone, although the functional consequences of these transcriptional changes remain to be determined.

## DISCUSSION

The present study demonstrates that pharmacological inhibition of HDACs can attenuate several consequences of mechanical unloading in skeletal muscle, providing further evidence that epigenetic regulation contributes to the development of disuse-associated muscle dysfunction. Using a murine HLS model, we found that TSA treatment partially preserved locomotor activity and was associated with improved muscle histological organization following 14 days of unloading. These functional and structural observations were accompanied by substantial changes in the skeletal muscle transcriptome, including attenuation of atrophy-associated gene expression, suppression of inflammatory and extracellular matrix remodeling programs, and restoration of several mitochondrial metabolic pathways. Importantly, TSA treatment was also associated with partial preservation of nuclear lamina architecture and H3K9me3-associated heterochromatin organization. Together, these findings suggest that HDAC-sensitive regulation influences multiple aspects of the skeletal muscle response to reduced mechanical loading, extending from tissue-level function to nuclear organization and transcriptional control.

Previous studies have established a role for HDACs in muscle wasting during disuse and denervation. In particular, Dupré-Aucouturier and colleagues^25^ demonstrated that TSA treatment attenuated soleus muscle atrophy during hindlimb suspension in rats, partially preserved myofiber size, and suppressed unloading-associated MuRF1 expression. Other studies have identified class I HDACs, particularly HDAC1, as regulators of FOXO activity and the muscle atrophy program^24^. Our findings are consistent with these observations, showing that HLS increased the expression of several atrophy-associated genes, including *Foxo1*, *Trim63*, *Fbxo32*, and *Cdkn1a*, whereas TSA treatment, particularly at the higher dose, shifted their expression toward baseline levels. We further observed altered expression of several HDAC family members, including *Hdac4* and *Hdac6*, suggesting that mechanical unloading produces a complex response involving multiple epigenetic regulators. Although these transcriptional changes do not establish the activity of individual HDAC isoforms, they support the involvement of HDAC-sensitive pathways in the regulation of muscle atrophy.

An important finding of this study is that the effects of TSA extend well beyond conventional muscle atrophy genes. Unloading produced widespread transcriptional changes, most prominently increased inflammatory and extracellular matrix remodeling signaling and suppressed mitochondrial metabolic pathways. TSA substantially reversed these patterns: high-dose TSA reversed many HLS-associated transcriptional signatures, compared with vehicle-treated mice. The DMSO vehicle produced modest improvements on its own, consistent with reports that DMSO has anti-inflammatory and antioxidant properties^26^. The TSA response was also not uniform across the transcriptome. Distinct gene modules differed in their sensitivity to TSA dose, and several mitochondrial pathways were enriched even relative to NS. These observations suggest that HDAC inhibition may not simply reverse unloading-induced changes but instead shift the muscle toward an alternative state with lower expression of catabolic and inflammatory genes and higher expression of mitochondrial energy metabolism programs. Whether these transcriptional changes translate into improved mitochondrial respiration, ATP production, or protein turnover remains to be established. A distinctive finding is the association between the transcriptional response to TSA and changes in nuclear architecture. Mechanical signals are transmitted to the nucleus through cytoskeletal connections, the LINC complex, and the nuclear lamina, which collectively contribute to the organization of chromatin and the regulation of gene expression. In our experiments, HLS was associated with irregular Lamin A/C distribution, altered nuclear morphology, and disruption of discrete H3K9me3-enriched heterochromatin foci. High-dose TSA partially preserved the peripheral organization of Lamin A/C and the focal distribution of H3K9me3. Interestingly, *Lmna* and *Syne1* transcript abundance increased following HLS despite the accompanying morphological abnormalities, raising the possibility that nuclear structural genes are transcriptionally regulated in response to altered mechanical loading. TSA treatment shifted several of these transcripts toward baseline levels while partially preserving nuclear architecture. These observations support a potential relationship between mechanical unloading, nuclear structural remodeling, and HDAC-sensitive chromatin regulation. However, altered H3K9me3 organization does not necessarily imply a corresponding change in total histone methylation or chromatin accessibility. Similarly, the present data do not directly demonstrate altered force transmission, nuclear mechanical properties, or lamina-associated chromatin interactions. Additional protein-level and quantitative imaging analyses will be necessary to establish how these nuclear structural changes relate to the transcriptional response.

An additional aspect of this study concerns the relationship between muscle preservation and bone adaptation during unloading. Although TSA treatment attenuated several muscle-associated consequences of HLS, it did not prevent the significant reduction in trabecular bone volume fraction observed after 14 days. Cortical bone architecture and whole-femur bending stiffness remained comparatively unchanged, suggesting that the early skeletal consequences of unloading were more apparent in trabecular bone than in the cortical compartment. Interestingly, muscle RNA-seq revealed altered expression of several candidate muscle-derived signaling factors associated with bone regulation, including *Ostn*, *Fndc5*, *Ccn4*, *Sfrp2*, and *Lcn2*, many of which shifted toward baseline following TSA treatment. These findings raise the possibility that HDAC inhibition modifies biochemical signaling potentially relevant to muscle–bone communication. However, changes in muscle transcript abundance do not establish altered secretion or biological activity of these factors. Moreover, systemic TSA administration could directly affect bone-resident cells, independently of its effects on skeletal muscle. The absence of trabecular protection despite substantial changes in muscle-associated signaling programs suggests that molecular responses in muscle and structural adaptations in bone are not necessarily coupled over the same timeframe.

Several limitations should be considered when interpreting these findings. The study examined a single unloading duration in female mice, and the functional measurements primarily reflect spontaneous locomotor activity rather than direct muscle contractile force. Although histology and ultrasound suggested partial preservation of muscle architecture, additional quantitative measurements of myofiber size and muscle mass would strengthen the assessment of structural protection. The transcriptomic analyses were performed using bulk muscle tissue, which contains multiple cellular populations, and therefore cannot distinguish transcriptional regulation within myofibers from changes in the abundance or activity of immune and stromal cells. Furthermore, although TSA was associated with substantial transcriptional rescue, the current experiments do not establish the specific HDAC isoforms or downstream protein-level mechanisms responsible. Immunoblotting to assess histone acetylation, H3K9me3 abundance, Lamin A/C, and proteins associated with FOXO-dependent atrophy pathways, together with quantitative immunofluorescence analysis of nuclear morphology and heterochromatin organization, will be important for strengthening the mechanistic interpretation. The systemic effects of high-dose TSA also warrant consideration, particularly the elevated serum transaminases observed in non-suspended mice. These observations highlight the need for further assessment of treatment safety, dosing, and tissue-specific effects before translational applications can be considered.

In summary, our findings indicate that HDAC inhibition attenuates multiple molecular and functional consequences of skeletal muscle unloading and is associated with partial preservation of nuclear lamina and heterochromatin organization. The results support a model in which mechanical deconditioning is accompanied by nuclear architectural changes and widespread transcriptional reprogramming, both of which are responsive to pharmacological HDAC inhibition. Further investigation of the causal relationships among nuclear mechanics, chromatin regulation, and muscle function may provide opportunities to develop more targeted approaches for mitigating disuse-induced muscle deterioration.

## ACKNOWLEDGEMENT

The authors acknowledge grant funding from NSF 2322878. The authors acknowledge Natalie Calahan for help in animal handling, Tia Tedford, Lauren Watts and Kelly Santangelo for help with histology, and Brad Nelson for help with ultrasound imaging. The authors utilized the ChatGPT for editing of the manuscript.

## SUPPLEMENTARY MATERIALS

**Supplementary Figure S1.**
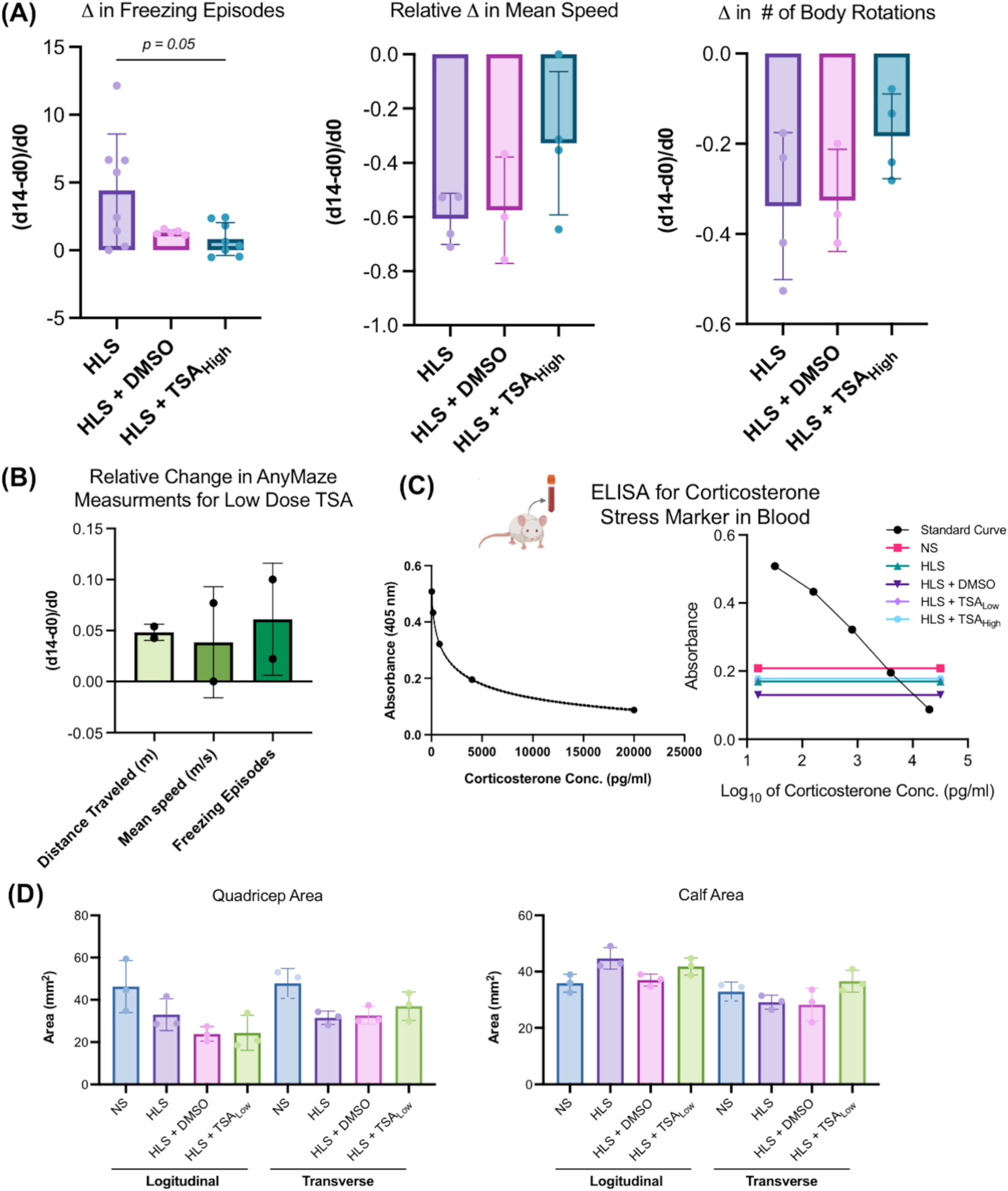
Additional analyses of locomotor activity, serum corticosterone, and hindlimb muscle morphology following hindlimb suspension and TSA treatment. **(A)** Quantification of additional locomotor parameters measured using ANY-maze before and after the 14-day hindlimb suspension period. Relative changes in freezing episodes, mean speed, and the number of body rotations are shown for HLS, HLS + DMSO, and HLS + high TSA groups. The increase in freezing episodes was reduced in the HLS + high TSA group compared with HLS (*p* = 0.05). Changes in mean speed and body rotations also showed trends toward improved locomotor activity following high-dose TSA treatment. Error bars represent mean ± SD, Kruskal–Wallis test followed by Dunn’s multiple comparisons test. **(B)** Relative changes in distance traveled, mean speed, and freezing episodes between baseline and day 14 in mice receiving HLS + low TSA (1.5 mg/kg body weight). Error bars represent mean ± SD. **(C)** ELISA standard curve used to quantify serum corticosterone concentrations. Absorbance at 405 nm is plotted against corticosterone concentration (left), together with the corresponding logarithmic standard curve and measured absorbance values for NS, HLS, HLS + DMSO, HLS + low TSA, and HLS + high TSA samples (right). **(D)** Quantification of quadriceps (thigh) and calf muscle areas from longitudinal and transverse ultrasound images acquired after 14 days of hindlimb suspension. Measurements are shown for NS, HLS, HLS + DMSO, and HLS + low TSA groups. Individual data points indicate measurements from individual mice. Data is not normalized to body weight. Error bars represent mean ± SD.

**Supplementary Figure S2.**
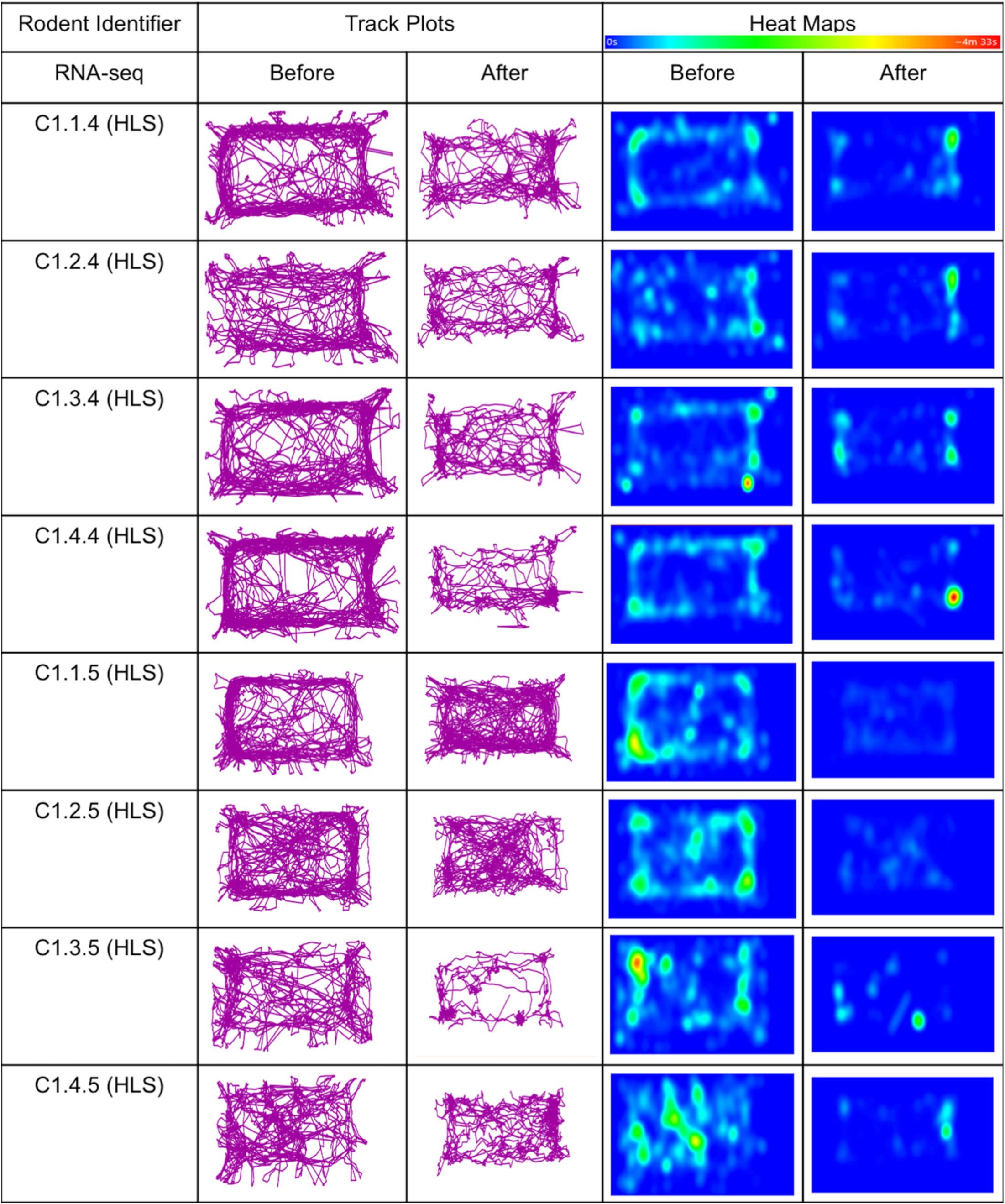
Individual locomotor trajectories and spatial occupancy heat maps of HLS mice. ANY-maze movement trajectories and corresponding spatial occupancy heat maps for individual mice before and after 14 days of hindlimb suspension (HLS). Heat maps represent the time spent in different regions of the arena, with warmer colors indicating longer occupancy.

**Supplementary Figure S3.**
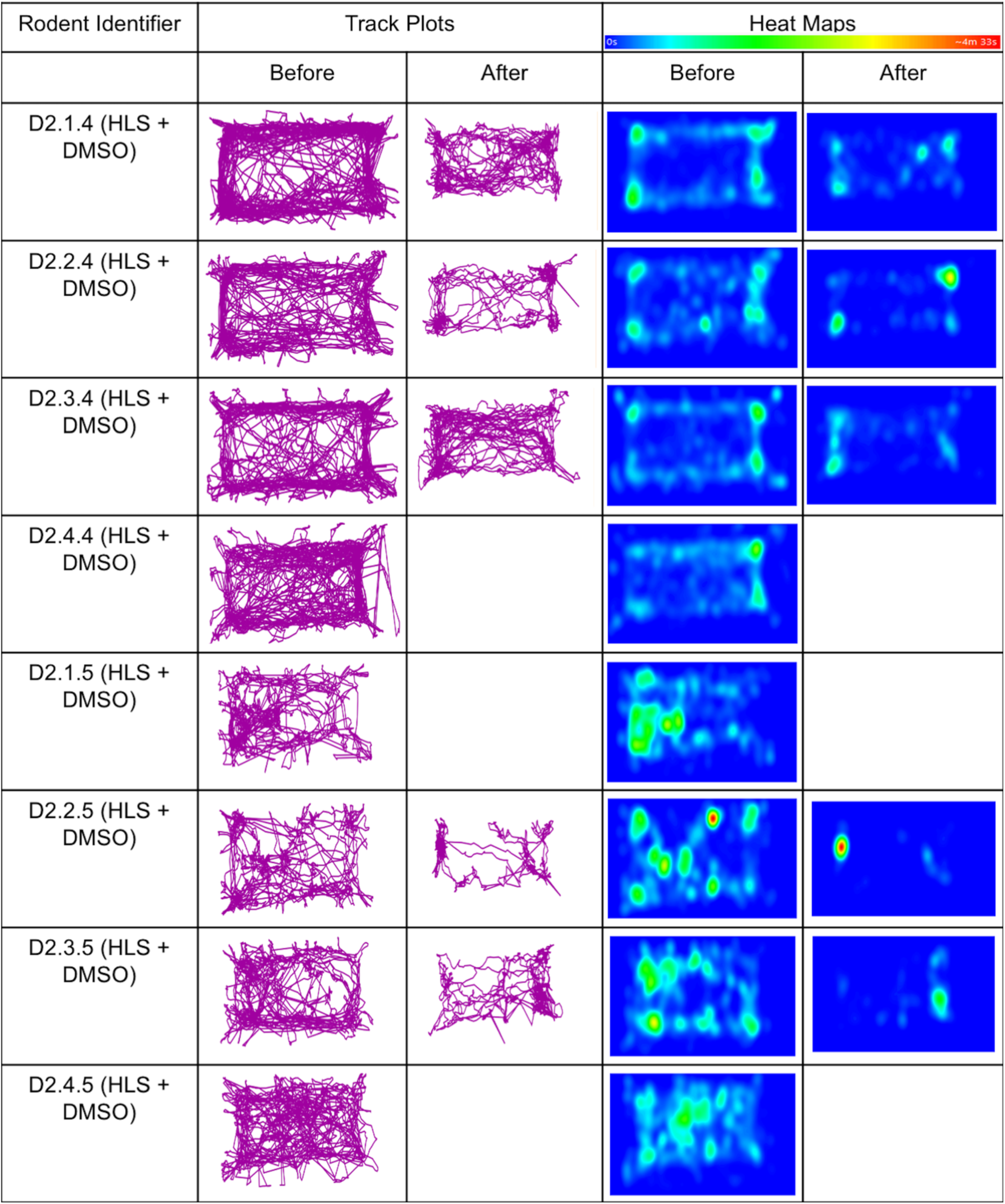
Individual locomotor trajectories and spatial occupancy heat maps of HLS + DMSO mice. ANY-maze movement trajectories and corresponding spatial occupancy heat maps for individual mice in the HLS + DMSO (vehicle) group before and after 14 days of hindlimb suspension. Heat maps indicate the spatial distribution of time spent within the arena. Day 14 (after) data was not captured for some mice that did not survive until the timepoint.

**Supplementary Figure S4.**
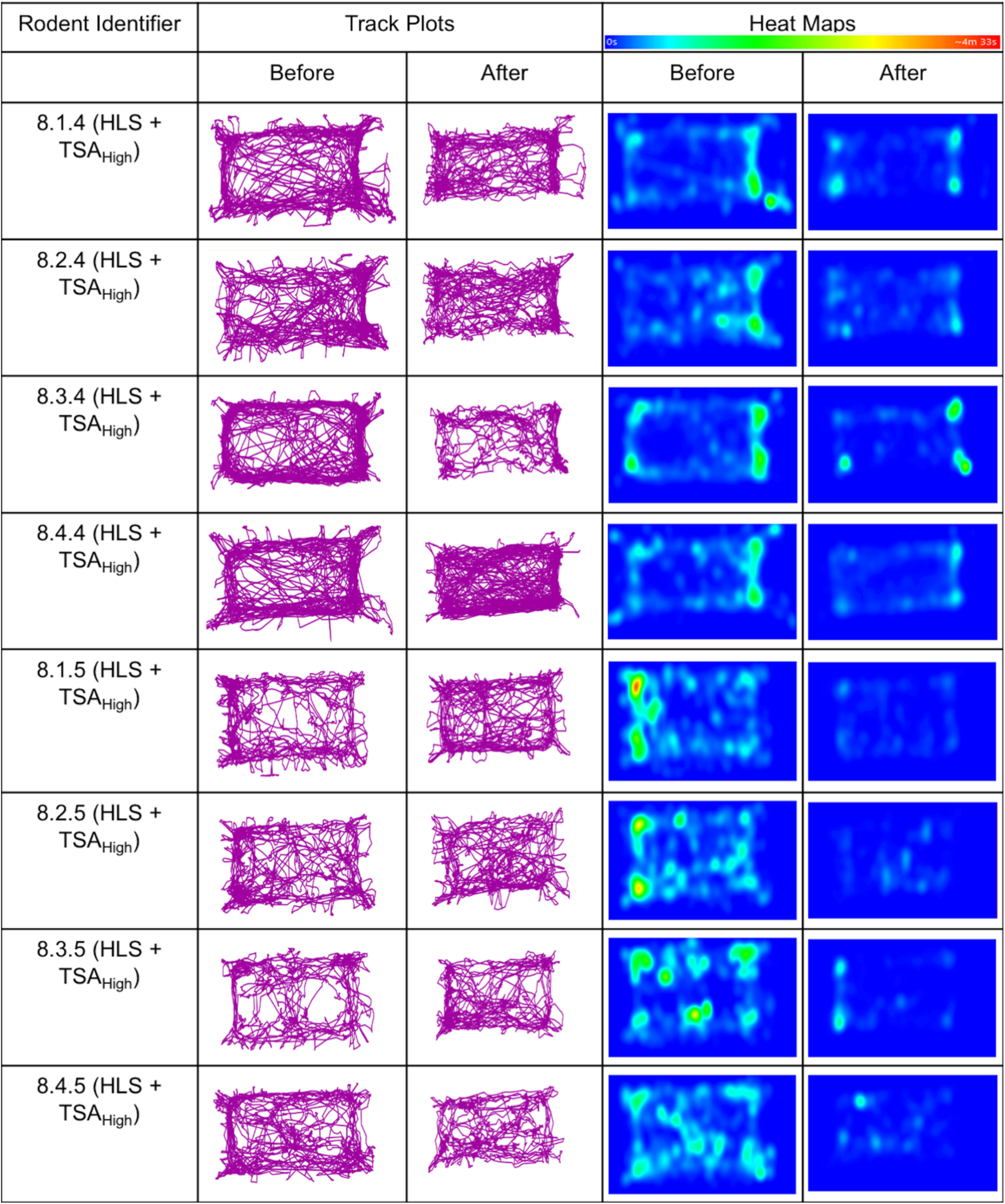
Individual locomotor trajectories and spatial occupancy heat maps of HLS + high TSA mice. ANY-maze movement trajectories and corresponding spatial occupancy heat maps for individual mice treated with high-dose TSA (8 mg/kg) before and after 14 days of hindlimb suspension. Movement trajectories illustrate the locomotor activity retained following TSA treatment.

**Supplementary Figure S5.**
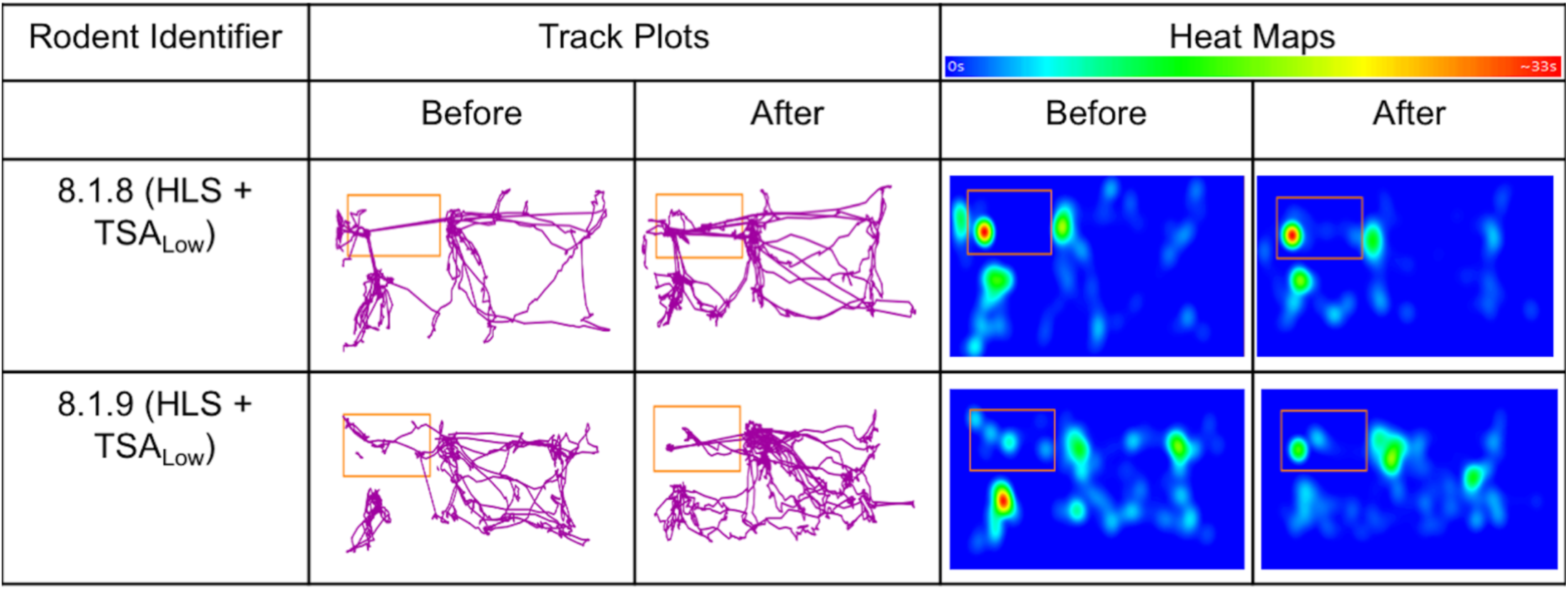
Individual locomotor trajectories and spatial occupancy heat maps of HLS + low TSA mice. ANY-maze movement trajectories and corresponding spatial occupancy heat maps for two mice treated with low-dose TSA (1.5 mg/kg) before and after 14 days of hindlimb suspension. These recordings are presented for completeness but were excluded from the primary locomotor analysis due to the limited sample size (n = 2).

**Supplementary Figure S6.**
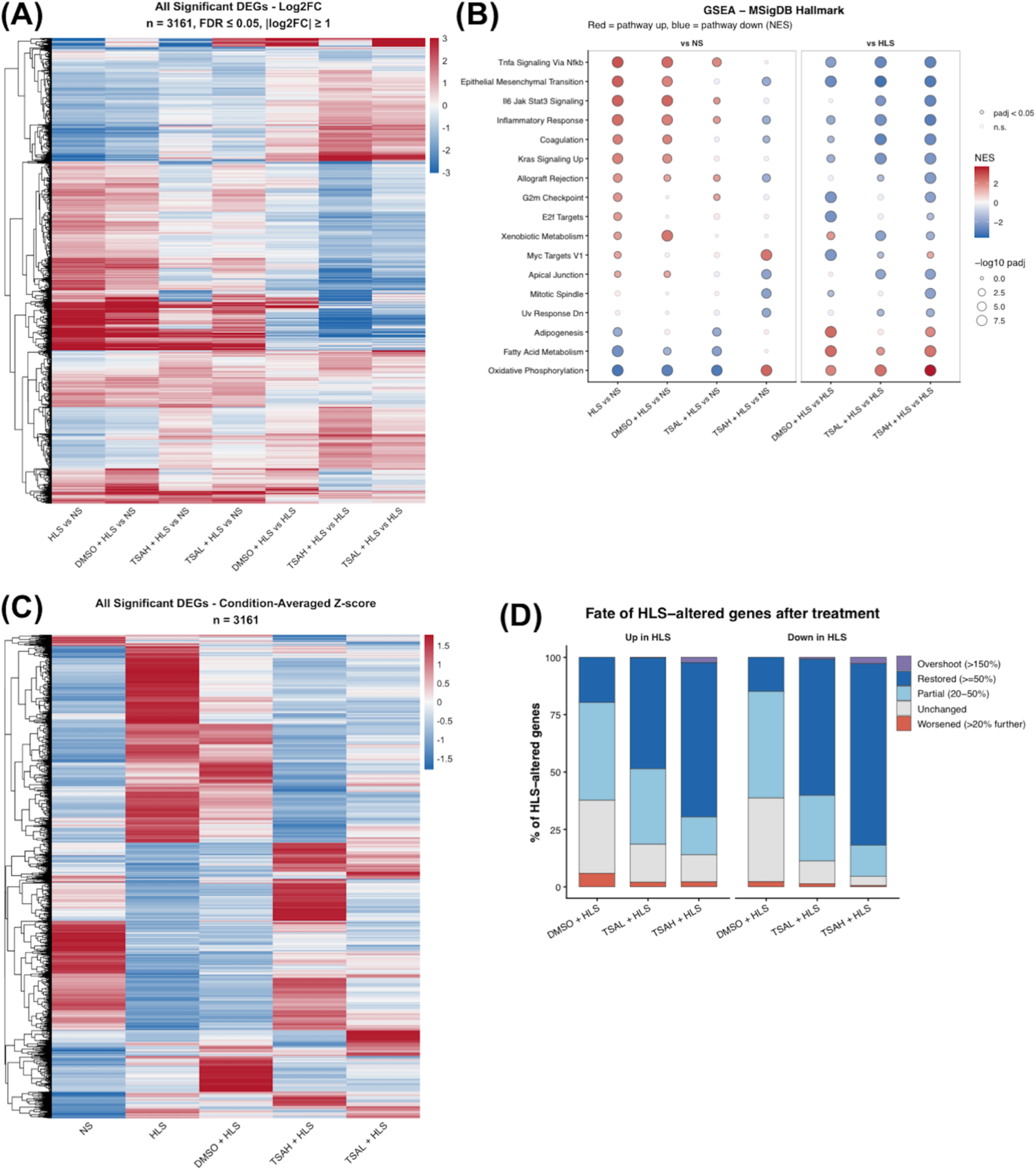
Transcriptomic analysis reveals broad reversal of unloading-associated gene expression changes and pathway enrichment following TSA treatment. **(A)** Heatmap of log2 fold changes for 3,161 differentially expressed genes identified across the indicated pairwise comparisons (FDR ≤ 0.05, |log2FC| ≥ 1). Red and blue indicate positive and negative fold changes, respectively, with genes organized by hierarchical clustering. **(B)** Gene set enrichment analysis (GSEA) using MSigDB Hallmark gene sets for comparisons relative to NS and HLS. Dot color represents the normalized enrichment score (NES), and dot size indicates the −log10 adjusted *p*-value. TSA treatment attenuated unloading-associated inflammatory signatures and increased the enrichment of oxidative phosphorylation pathways. **(C)** Heatmap showing condition-averaged Z-scores for the 3,161 DEGs across NS, HLS, HLS + DMSO, HLS + high TSA, and HLS + low TSA groups, illustrating treatment-associated changes in global expression patterns. **(D)** Classification of HLS-altered genes according to their response to treatment. Genes initially upregulated or downregulated by HLS were categorized as restored (≥50% reversal), partially restored (20–50%), unchanged, worsened, or overshooting the baseline expression level. High-dose TSA produced the greatest proportion of restored genes in both categories, compared with low-dose TSA and DMSO treatment.

**Supplementary Figure S7.**
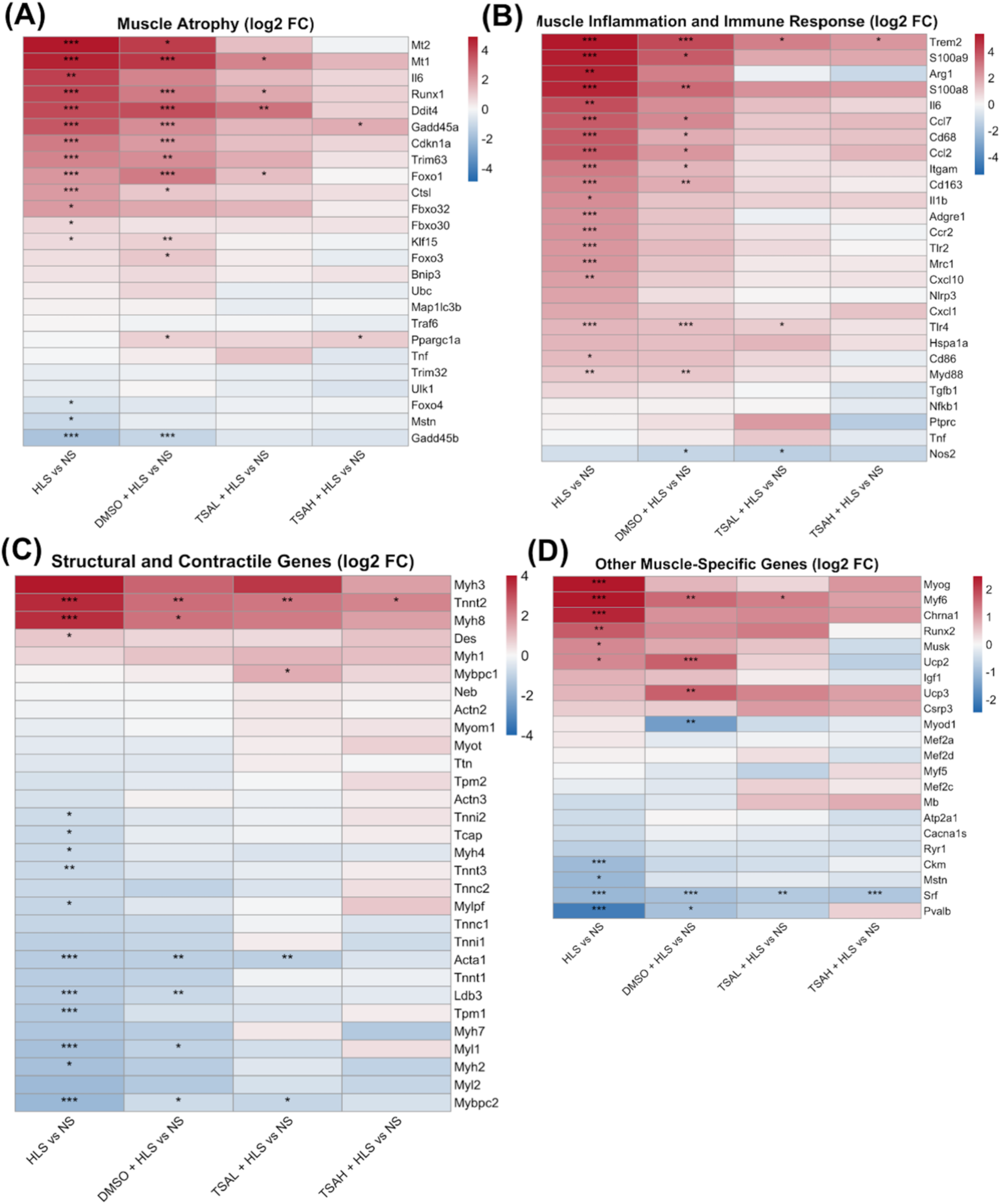
TSA treatment modifies unloading-associated transcriptional changes in muscle atrophy, inflammation, and structural gene programs. Heatmaps showing RNA-seq-derived log2 fold changes relative to NS for selected genes associated with **(A)** muscle atrophy and protein degradation, **(B)** inflammation and immune responses, **(C)** muscle structural and contractile proteins, and **(D)** other muscle-associated regulatory and functional processes. Columns represent HLS, HLS + DMSO, HLS + low TSA, and HLS + high TSA, each compared with NS. Red and blue indicate increased and decreased gene expression, respectively. Statistical significance was determined using DESeq2-adjusted *p*-values: * adjusted *p* < 0.05, ** adjusted *p* < 0.01, *** adjusted *p* < 0.001.

**Supplementary Figure S8.**
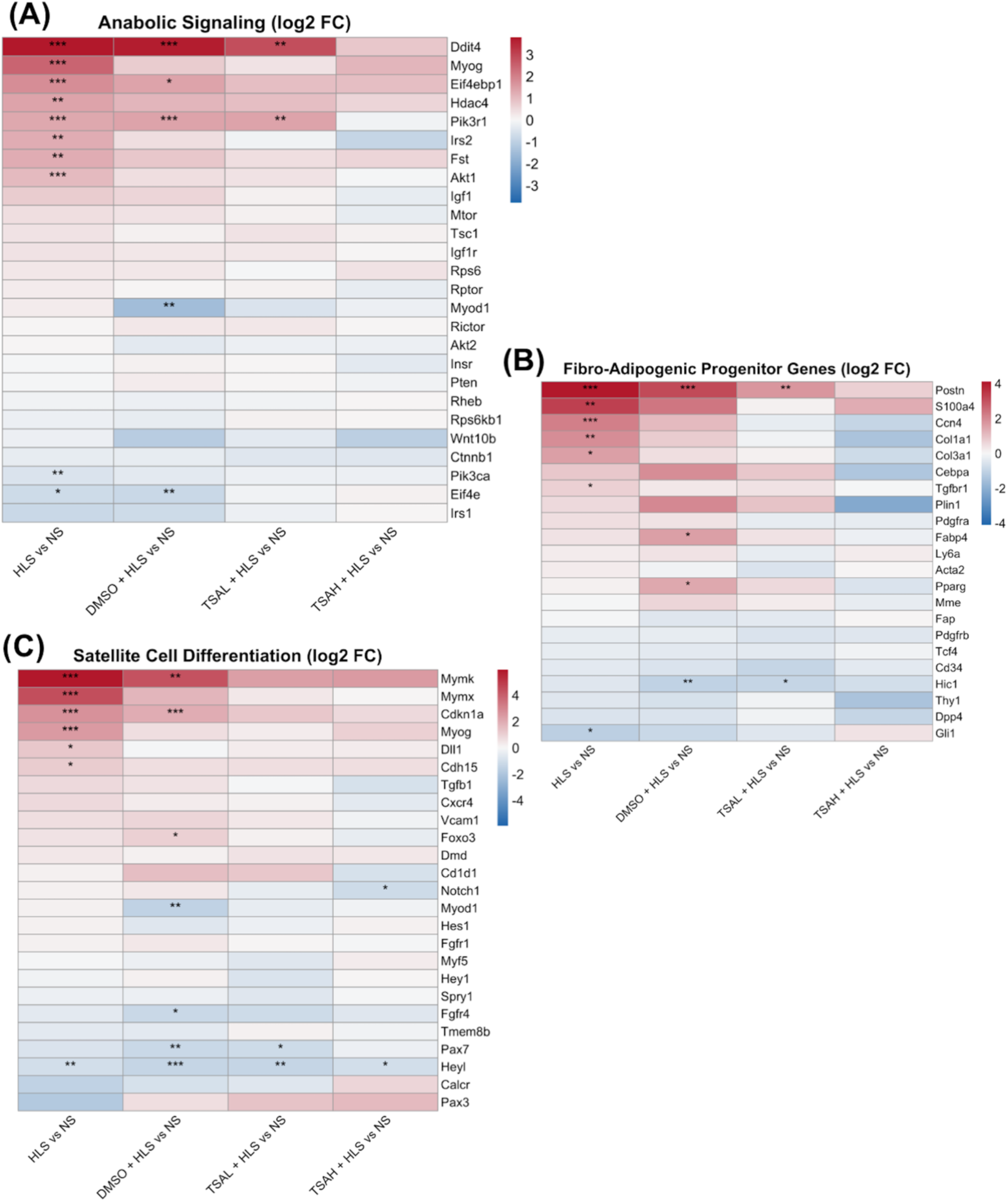
Transcriptional changes in anabolic signaling, fibro-adipogenic progenitor-associated genes, and muscle satellite cell differentiation following TSA treatment. Heatmaps showing log2 fold changes relative to NS for selected genes associated with **(A)** anabolic signaling and its regulation, **(B)** fibro-adipogenic progenitor-associated signaling and extracellular matrix remodeling, and **(C)** satellite cell differentiation and myogenic regulation. Expression profiles are shown for HLS, HLS + DMSO, HLS + low TSA, and HLS + high TSA relative to NS. Red and blue indicate increased and decreased expression, respectively. Asterisks indicate DESeq2-adjusted statistical significance: * adjusted *p* < 0.05, ** adjusted *p* < 0.01, *** adjusted *p* < 0.001.

**Supplementary Figure S9.**
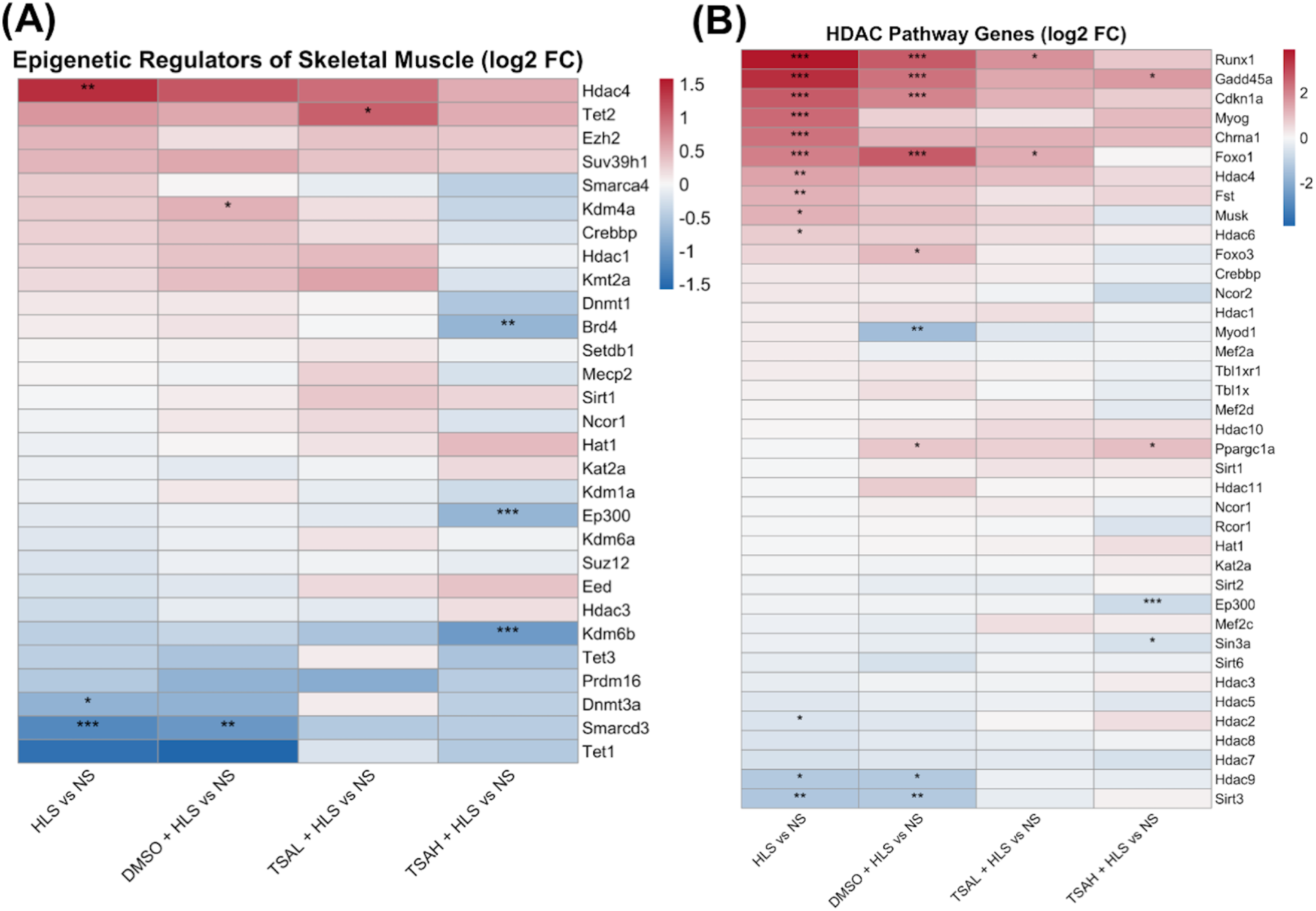
HDAC inhibition alters the transcriptional profiles of epigenetic regulators and HDAC-associated genes following hindlimb suspension. Heatmaps showing log2 fold changes relative to NS for **(A)** selected epigenetic regulators involved in chromatin organization, histone modification, and DNA methylation, and **(B)** HDAC family members and genes associated with HDAC-related transcriptional regulation. Expression changes are presented for HLS, HLS + DMSO, HLS + low TSA, and HLS + high TSA relative to NS. Red and blue indicate increased and decreased expression, respectively. Asterisks denote DESeq2-adjusted statistical significance: * adjusted *p* < 0.05, ** adjusted *p* < 0.01, *** adjusted *p* < 0.001.

**Supplementary Figure S10.**
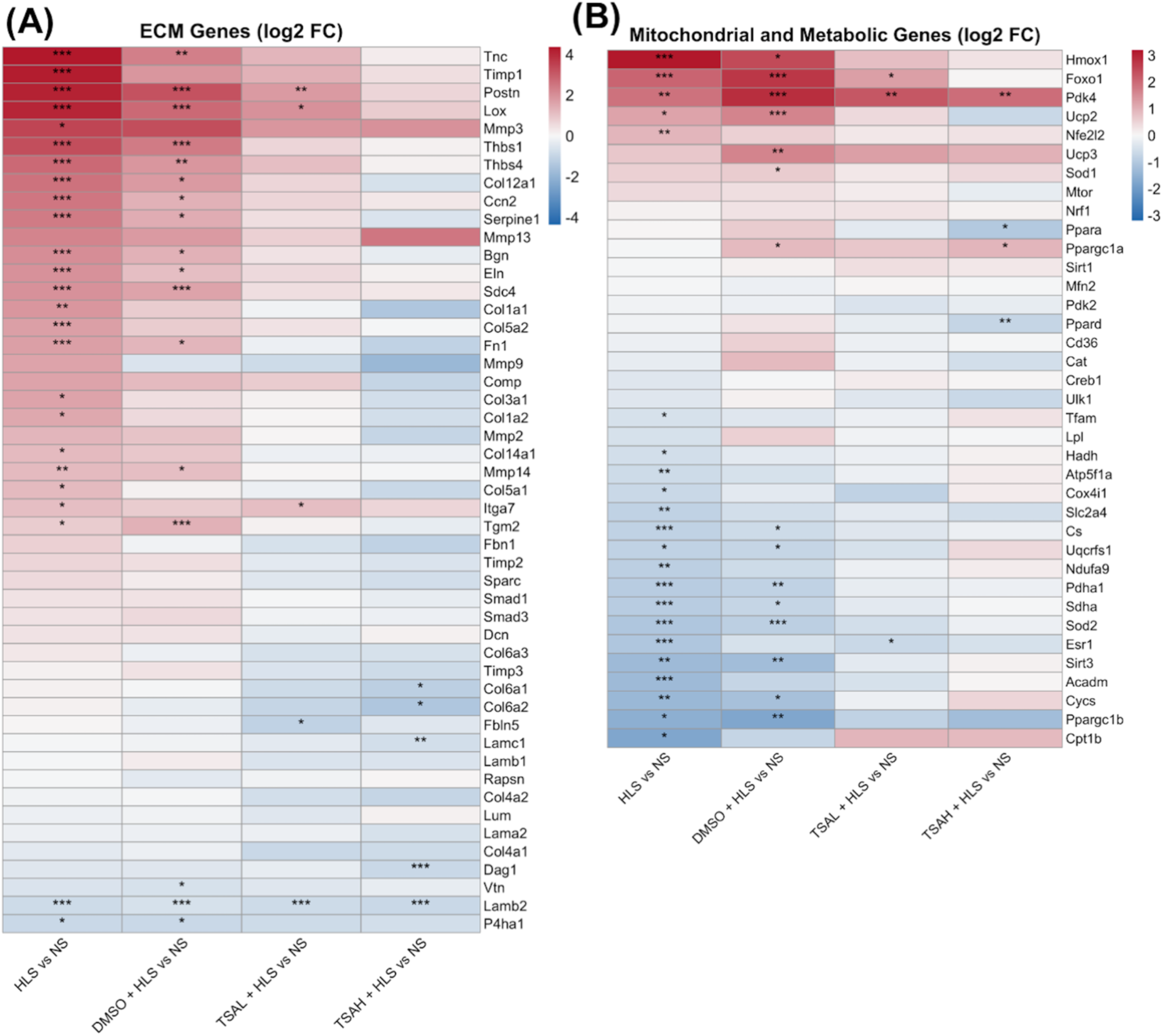
TSA treatment modifies extracellular matrix and mitochondrial metabolism-associated transcriptional programs following hindlimb suspension. Heatmaps showing log2 fold changes relative to NS for selected genes associated with **(A)** extracellular matrix organization, collagen regulation, cell adhesion, and matrix remodeling, and **(B)** mitochondrial function, energy metabolism, and oxidative stress responses. HLS-associated transcriptional changes were broadly attenuated following TSA treatment, with distinct responses among individual genes and doses. Columns represent HLS, HLS + DMSO, HLS + low TSA, and HLS + high TSA relative to NS. Asterisks indicate DESeq2-adjusted statistical significance: * adjusted *p* < 0.05, ** adjusted *p* < 0.01, *** adjusted *p* < 0.001.

**Supplementary Figure S11.**
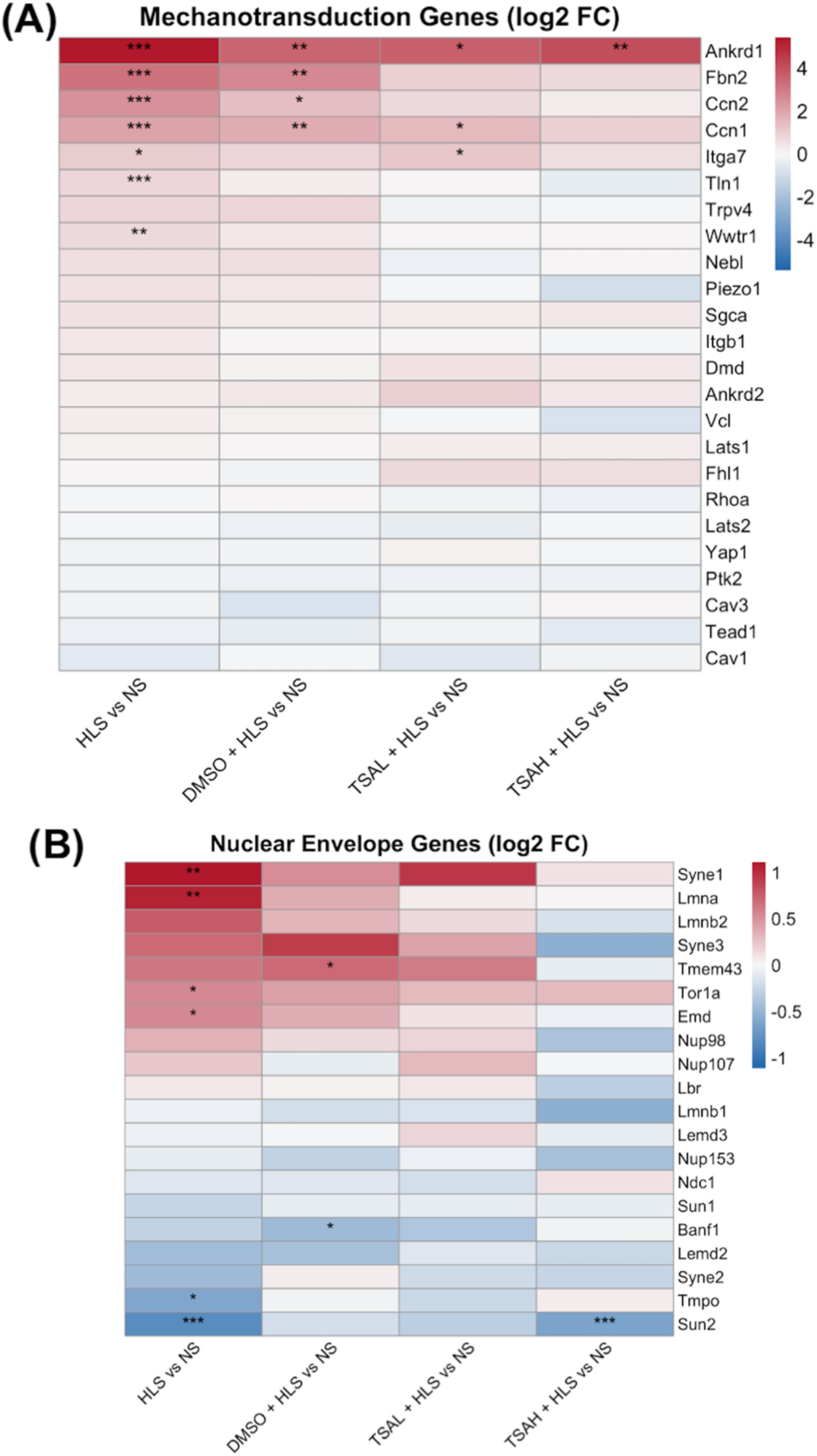
Transcriptional responses of mechanotransduction and nuclear envelope-associated genes to mechanical unloading and TSA treatment. Heatmaps showing log2 fold changes relative to NS for **(A)** genes involved in mechanotransduction, cytoskeletal regulation, and cell–matrix signaling, and **(B)** genes encoding nuclear envelope, nuclear lamina, and nuclear– cytoskeletal coupling proteins. Expression patterns are shown for HLS, HLS + DMSO, HLS + low TSA, and HLS + high TSA relative to NS. Red and blue indicate increased and decreased transcript abundance, respectively. Asterisks indicate DESeq2-adjusted statistical significance: * adjusted *p* < 0.05, ** adjusted *p* < 0.01, *** adjusted *p* < 0.001.

**Supplementary Figure S12.**
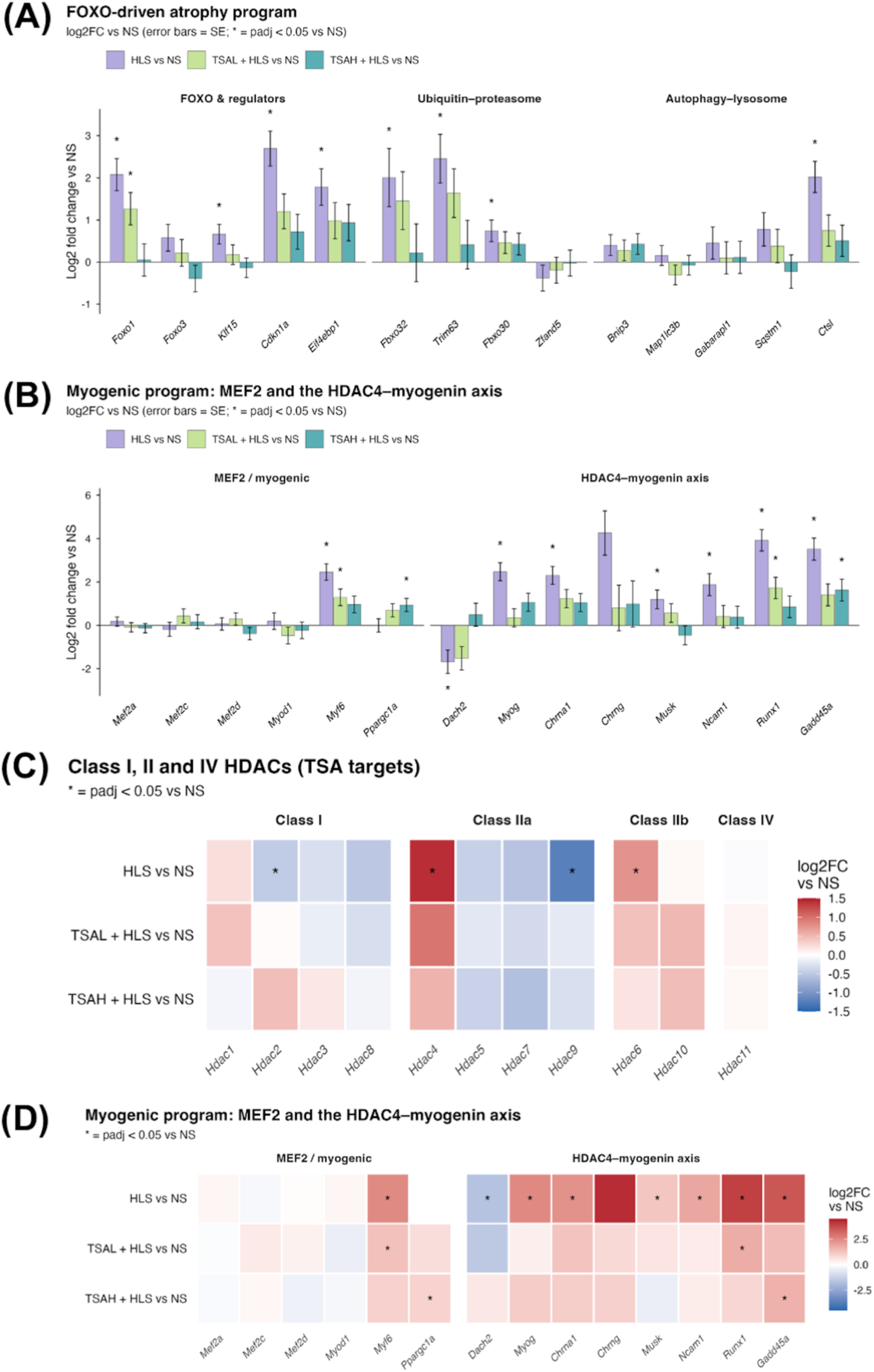
Detailed transcriptional analysis of FOXO-associated atrophy, myogenic regulation, and HDAC family genes following TSA treatment. **(A)** RNA-seq-derived log2 fold changes relative to NS for selected genes involved in FOXO-associated regulation, ubiquitin–proteasome-mediated protein degradation, and autophagy–lysosome pathways. **(B)** Log2 fold changes for genes associated with MEF2-dependent myogenic regulation and HDAC4-related muscle remodeling, including *Myf6, Myog, Chrna1, Runx1,* and *Gadd45a*. Error bars in (A) and (B) represent standard error. **(C)** Heatmap showing log2 fold changes of Class I, II, and IV HDAC family transcripts relative to NS. **(D)** Heatmap showing expression changes in MEF2-associated myogenic and muscle remodeling genes. All comparisons are relative to NS and include HLS, HLS + low TSA, and HLS + high TSA groups. Asterisks indicate DESeq2-adjusted *p* < 0.05.

**Supplementary Figure S13.**
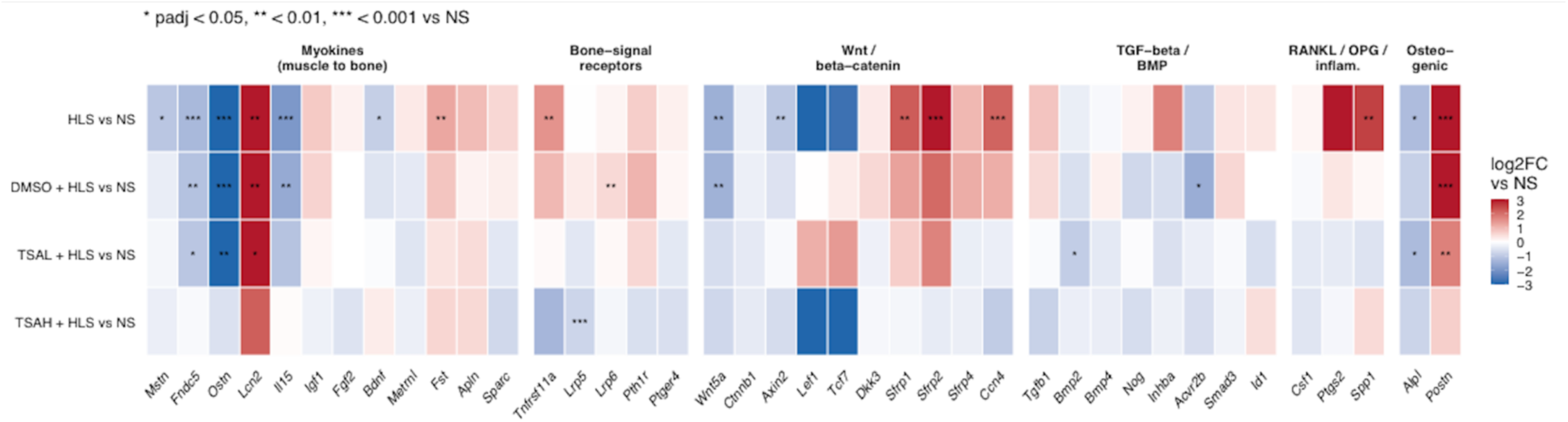
Transcriptional changes in muscle-associated signaling factors and pathways potentially involved in muscle-bone communication following hindlimb suspension and TSA treatment. Heatmap showing RNA-seq-derived log2 fold changes relative to NS for selected genes grouped according to their involvement in muscle-derived signaling (putative myokines), bone-associated signaling receptors, Wnt/β-catenin regulation, TGF-β/BMP signaling, RANKL/OPG-associated inflammatory regulation, and osteogenic processes. Expression changes are shown for HLS, HLS + DMSO, HLS + low TSA, and HLS + high TSA relative to NS. Mechanical unloading altered the expression of several muscle-associated signaling factors and regulatory genes, with heterogeneous responses following TSA treatment. Red and blue indicate increased and decreased transcript abundance, respectively. Asterisks indicate DESeq2-adjusted statistical significance relative to NS: * adjusted *p* < 0.05, ** adjusted *p* < 0.01, *** adjusted *p* < 0.001.

## REFERENCES

1. Bodine, S. C. Disuse-induced muscle wasting. International Journal of Biochemistry & Cell Biology (2013).

2. Sartori, R., Romanello, V. & Sandri, M. Mechanisms of muscle atrophy and hypertrophy: implications in health and disease. Nature Communications 1–12 (2021).

3. Brooks, N. E. & Myburgh, K. H. Skeletal muscle wasting with disuse atrophy is multi-dimensional : the response and interaction of myonuclei, satellite cells and signaling pathways. Frontiers in Physiology 5, 1–14 (2014).

4. Larsson, X. L. et al. Sarcopenia: aging related loss of muscle mass and function. Physiological Reviews 427–511 (2025).

5. Argiles, J. M., Lopez-Soriano, F. J., Stemmler, B. & Busquets, S. Cancer-associated cachexia — understanding the tumour macroenvironment and microenvironment to improve management. Nature Reviews Clinical Oncology (2023).

6. Gao, Y., Arfat, Y., Wang, H. & Goswami, N. Muscle atrophy induced by mechanical unloading : Mechanisms and potential countermeasures. Frontiers in Pysiology 9, (2018).

7. Vandenburgh, H., Chormiak, J., Shansky, J., Del Tatto, M. & Lemarie, J. Space travel directly induces skeletal muscle atrophy. FASEB Journal 13, 1031–8 (1999).

8. Kortebien, P., Symons, T. B. & Ferrando, A. Functional impact of 10 days of bed rest in healthy older adults. J Gerontol A Biol Sci Med Sci (2008).

9. English, K. L. & Paddon-Jones, D. Protecting muscle mass and function in older adults during bed rest. Current Opinion in Clinical Nutrition and Metabolic Care (2010).

10. Musaro, A. Muscle homeostasis and regeneration: From molecular mechanisms to therapeutic opportunities. Cells (2020).

11. Gehlert, S., Bloch, W. & Suhr, F. Ca dependent regulations and signaling in skeletal muscle : from electro-mechanical coupling to adaptation. International Journal of Molecular Sciences 1066–1095 (2015).

12. Tidball, J. G. Mechanical signal transduction in skeletal muscle growth and adaptation. Journal of Applied Physiology 1900–1908 (2025).

13. Glass, D. J. Signalling pathways that mediate skeletal muscle hypertrophy and atrophy. Nature Cell Biology (2003).

14. Bodine, S. C. & Baehr, L. M. Skeletal muscle atrophy and the E3 ubiquitin ligases MuRF1 and MAFbx/atrogin-1. Am J Physiol Endocrinol Metab (2014).

15. Orr, A. W., Helmke, B. P., Blackman, B. R. & Schwartz, M. A. Mechanisms of mechanotransduction. Developmental Cell 10, 11–20 (2006).

16. Iskratsch, T., Wolfenson, H. & Sheetz, M. P. Appreciating force and shape - the rise of mechanotransduction in cell biology. Nature Reviews Molecular Cell Biology 15, 825–33 (2014).

17. Wang, N., Tytell, J. D. & Ingber, D. E. Mechanotransduction at a distance: mechanically coupling the extracellular matrix with the nucleus. Nature Reviews Molecular Cell Biology 10, 75–82 (2009).

18. Isermann, P. & Lammerding, J. Nuclear mechanics and mechanotransduction in health and disease. Current Biology 23, R1113–R1121 (2013).

19. Méjat, A. & Misteli, T. LINC complexes in health and disease. Nucleus 1, 40–52 (2010).

20. Seelbinder, B. et al. Nuclear deformation guides chromatin reorganization in cardiac development and disease. Nature Biomedical Engineering 5, (2021).

21. Seaborne, R. A. et al. Human skeletal muscle possesses an epigenetic memory of hypertrophy. Scientific Reports 1–17 (2018).

22. Sharples, A. P., Stewart, C. E. & Seaborne, R. A. Does skeletal muscle have an ‘epi’-memory? The role of epigenetics in nutritional programming, metabolic disease, aging and exercise. Aging Cell 603–616 (2016).

23. Bongers, K. S. et al. Skeletal muscle denervation causes skeletal muscle atrophy through a pathway that involves both Gadd45a and HDAC4. American Journal of Physiology-Endocrinology and Met 305, (2013).

24. Beharry, A. W. et al. HDAC1 activates FoxO and is both sufficient and required for skeletal muscle atrophy. Journal of Cell Science 127, 1441–1453 (2014).

25. Dupre-Aucouturier, S., Castells, J., Freyssenet, D. & Desplanches, D. Trichostatin A, a histone deacetylase inhibitor, modulates unloaded-induced skeletal muscle atrophy. Journal of Applied Physiology 119, 342–351 (2015).

26. Santos, N. C., Figueira-Coelho, J., Martins-Silva, J. & Saldanha, C. Multidisciplinary utilization of dimethyl sulfoxide: pharmacological, cellular, and molecular aspects. Biochemical Pharmacology 65, 1035–1041 (2003).

